# Comparative study of Ergosterol and 7-dehydrocholesterol and their Endoperoxides: Generation, Identification and Impact in Phospholipid Membranes and Melanoma Cells

**DOI:** 10.1101/2024.10.21.619400

**Authors:** Megumi Nishitani Yukuyama, Karen Campos Fabiano, Alex Inague, Miriam Uemi, Rodrigo Santiago Lima, Larissa Regina Diniz, Tiago Eugenio Oliveira, Thais Satie Iijima, Hector Oreliana Fernandes Faria, Rosangela Silva Santos, Maria Fernanda Valente Nolf, Adriano Brito Chaves-Filho, Marcos Yukio Yoshinaga, Helena Couto Junqueira, Paolo Di Mascio, Mauricio da Silva Baptista, Sayuri Miyamoto

## Abstract

Melanoma is an aggressive cancer that has attracted attention in recent years due to its high mortality rate of 80%. Damage caused by oxidative stress generated by radical (type I reaction) and singlet oxygen, ^1^O_2_ (type II reaction) oxidative reactions may induce cancer. Thus, studies that aim to unveil the mechanism that drives these oxidative damage processes become relevant. Ergosterol, an analogue of 7-dehydrocholesterol, important in the structure of cell membranes, is widely explored in cancer treatment. However, to date little is known about the impact of different oxidative reactions on these sterols in melanoma treatment, and conflicting results about their effectiveness complicates the understanding of their role in oxidative damage. Our results highlight differences among ergosterol, 7-dehydrocholesterol (7-DHC) and cholesterol in membrane properties when subjected to distinct oxidative reactions. Furthermore, we conducted a comparative study exploring the mechanisms of cell damage by photodynamic treatment in A375 melanoma. Notably, endoperoxides from ergosterol and 7-DHC generated by ^1^O_2_ showed superior efficacy in reducing the viability of A375 cells compared to their precursor molecules. We also describe a step-by-step process to produce and identify endoperoxides derived from ergosterol and 7-DHC. While further studies are needed, this work provides new insights for understanding cancer cell death induced by different oxidative reactions in the presence of biologically relevant sterols.

## INTRODUCTION

Skin cancer was reported to be the fifth most common cancer worldwide in 2020. It is divided in non-melanoma (basal and squamous cell carcinoma) and melanoma. Although melanoma corresponds to only 4% of total skin cancer, it is considered the most dangerous due to its high death rate of 80% (1). A total of 325,000 new cases of melanoma and 57,000 deaths were estimated for 2020 worldwide, and an increase of approximately 50% of new cases and 68% deaths are projected for 2040 (2). Thus, a convenient treatment becomes crucial for improving the survival rate of this disease.

Sunlight exerts many beneficial effects such as vitamin D synthesis, anti-inflammatory stimulation, and prevention against several diseases (e.g., certain types of cancers, diabetes, and multiple sclerosis) (3). However, excess radiation may lead to negative effects causing oxidative stress and inflammation in skin cells: both UV and visible light have the potential to promote photoexcitation of endogenous photosensitizers. In this process light energy is converted into chemical energy producing reactive oxygen species (ROS), thereby leading to redox imbalance in the cells under excessive radiation (4). Examples of endogenous photosensitizers are porphyrins, flavin, melanin, and lipofuscin, and the concentration and presence of these photosensitizers influence the sensitivity of specific tissue or cells to photoinduced damage (5). Photooxidation reactions are classified as Type I or Type II reactions. Type I reaction generates reactive radical species, such as superoxide radical anion (O_2_ ^•−^) and hydroperoxyl radicals (HOO^-^), while type II reaction generates singlet molecular oxygen (^1^O_2_) (6). Although current photoprotection products block UV rays, emerging studies show that skin photodamage by visible light is responsible for more than 50% of radicals generated under sun exposure (4, 7). In a recent review article, photoinduced damage on cellular DNA and skin was attributed to immediate and late oxidative reactions induced by UVA and visible light (7). The authors proposed that UVA radiation significantly contributes to the generation of 8-oxo-7,8-dihydroguanine through the excitation of endogenous photosensitizers, an immediate reaction mainly mediated by ^1^O_2_. In contrast, the late generation of reactive species is attributed to chemoexcitation of “dark” cyclobutane pyrimidine dimers. The article also highlights the influence of the blue component of visible light on cellular DNA damage (7).

Among several biomolecules, lipids play essential roles in cell membrane structure, serve as energy source, and generate metabolites that support proper cell development and proliferation (8) Cholesterol is an essential component of cell membranes and a precursor for other steroidal biomolecules such as hormones and vitamin D (9, 10). When oxidized, cholesterol gives rise to oxidized derivatives, known as oxysterols, including reactive sterol electrophiles that may disturb cellular function, leading to several diseases including cancer (11).

Ergosterol (Erg) was first discovered in 1889 in extracts of the ergot fungus (*Claviceps purpurea*) by Tanret and described as the major sterol in fungi (12). Likewise cholesterol, Erg is found mainly in the lipid bilayer of the cell membranes of most fungi, playing a crucial role in cell maintenance and growth (13). The additional double bonds at the C_7_ and C_22_ positions and a methyl bond at the C_24_ position differentiate Erg from cholesterol (**Figure 1**). Increasing attention has been given to Erg and its derivatives given their potential to inhibit cell growth efficacy in several cancer cells, by interacting with JAK/STAT mechanism or caspase-independent apoptosis (14, 15). Additionally, studies have indicated that oxidized derivatives of Erg, such as the 5α, 8α-epidioxiergosta-6 and 22-dien-3β-ol (EEP), induce human hepatocellular carcinoma cell death through the activation of pro-apoptotic protein Puma and Bax under stimulation of Foxo3 activity (16).

**Figure 1.**
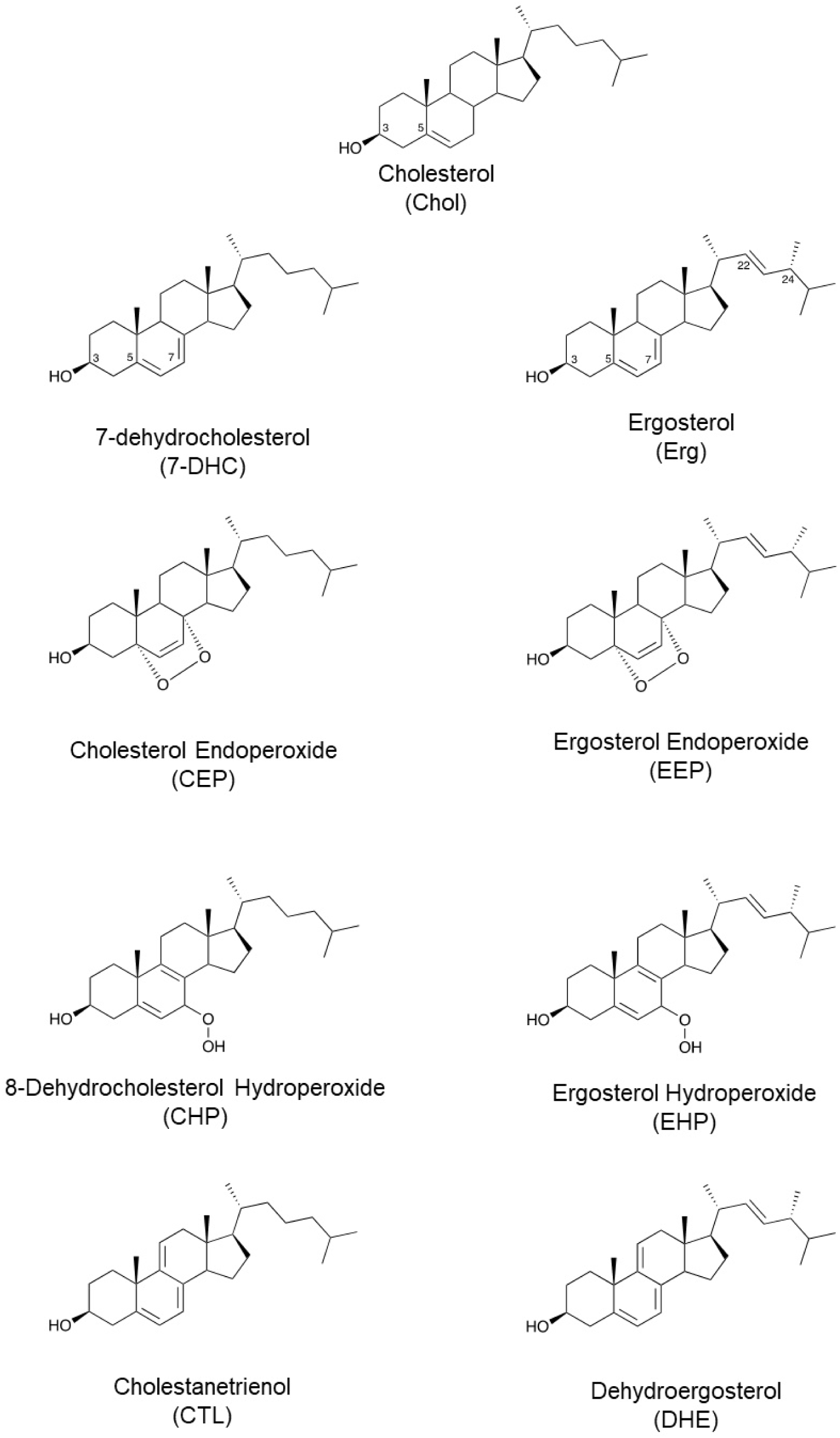
Structure of 7-dehydrocholesterol, ergosterol and their oxidation products.

7-dehydrocholesterol (7-DHC) is a precursor of cholesterol in mammals, with a very similar structure to Erg (**Figure 1**) (17, 18). The anticancer effect of 7-DHC has been demonstrated by the dose-dependent decrease in viability of melanoma A2058 cell line. In the absence of light, cells treated with 7-DHC generated ROS, increased pro-apoptotic Bax protein, and affected mitochondrial permeabilization by reducing mitochondria membrane potential (19). Additionally, in ovarian cancer cell lines, replacing cholesterol with 7-DHC significantly enhanced the anticancer activity of photosensitizer-encapsulated liposomes upon irradiation (20). In contrast, other studies demonstrated that 7-DHC suppressed and regulated cell death by ferroptosis, an iron-dependent non-apoptotic cell death caused by exacerbated lipid peroxidation (21, 22). For instance, it has been shown that pre-treatment of B16F10 melanoma cells with 7-DHC significantly protected cells from ferroptosis, suggesting that 7-DHC can increase melanoma resistance against oxidative stress (21).

Given the conflicting roles of Erg and 7-DHC in cancer treatment, we sought to compare the damage caused by these sterols and their respective endoperoxides under oxidation reactions type I and type II in mimetic lipid membranes and cancer cells. Herein, we present a step-by-step process to produce endoperoxide derivatives of Erg and 7-DHC and evaluate they role in lipid bilayers and melanoma A375 cells under Type I or Type II photooxidative damages.

## MATERIALS

Phospholipids palmitoyl-oleoyl phosphatidylcholine (POPC, 1-palmitoyl-2-oleoyl-sn-glycero-3-phosphocholine) and egg yolk phosphatidylcholine (EggPC, L-α-phosphatidylcholine) were purchased from Avanti Polar Lipids (Alabaster, USA). Solvents (HPLC grade) were purchased from Merck (Darmstadt, Germany), methylene blue, dimethy-methylene blue (DMMB) and other reagents were purchased from Sigma-Aldrich (St. Louis, USA). Fetal bovine serum, penicillin-streptomycin, and Dulbecco’s Modified Eagle’s Medium-high glucose (DMEM high glucose GlutaMAX, Supplement catalog number 10569010), and pyruvate were purchased from Gibco (ThermoFischer Scientific). Deuterium chloroform (CDCl_3_) was purchased from Sigma Aldrich (Steinheim, Germany).

## METHODS

### Synthesis and purification of sterol oxidation products

#### Sterol photo-oxidation

First, 200 mg of sterol (Erg, 7-DHC and cholesterol) were dissolved in 40 mL of chloroform in a 100 mL round-bottom flask, and 50 µL of the photosensitizer methylene blue (10 mM in methanol) was added. This solution was placed in an ice bath and an aliquot was collected at time zero. The mixture of sterol and methylene blue was kept under oxygen flow and stirred under a 500 W tungsten lamp for 90 minutes (Erg and 7-DHC) or 5 hours (cholesterol). Aliquots were collected in every 30 minutes for qualitative analysis by thin layer chromatography (TLC) using a silica plate eluted with ethyl acetate and isooctane (1:1 v/v). Plates were air-dried, sprayed with 50% sulfuric acid, and heated to allow visualization of sterols and their oxidized products. (**Figure S1).**

#### Purification of sterol oxidation products

Sterol photooxidation products were initially separated by flash column chromatography containing silica gel in hexane. After equilibrating the column with hexane, a gradient of hexane and ethyl acetate in different volume ratios, 95:5 (50 mL), 90:10 (50 ml), 80:20 (100 ml), 70:30 (100 ml), 60:40 (100 mL), 50:50 (50 ml) and 40:60 (50 ml) was used for elution. Fractions of approximately 10 mL were collected and checked by TLC (**Figure S2**). Sterol oxidized products were further purified by semi-preparative HPLC (LC 20AD, Prominence, Shimadzu, Kyoto, Japan) coupled to a Diode Array Detector (DAD, SPDM20A, Prominence, Shimadzu, Kyoto, Japan) using the following conditions: Luna C18 column (250 x 10 mm, 5 µm, 100 Å); flow rate 5 mL/min; 40°C and mobile phase consisted of 95% methanol and 5% water. Each peak was collected in an amber glass tube using a fraction collector (FRC10-A, Shimadzu, Kyoto, Japan) and dried with the rotary evaporator.

### Characterization of sterol photooxidation products

#### Analysis of oxidation products by HPLC-DAD

Sterol oxidation products were analyzed by HPLC (Prominence, Shimadzu, Kyoto, Japan) coupled to a diode array detector (DAD) using a Gemini C18 column (250 x 4.6 mm; 5 µm, 110 Å), in isocratic elution at a flow rate of 1 mL/min with a mixture of 95 % methanol and 5% water as mobile phase.

#### Analysis of oxidation products by mass spectrometry

The analysis of oxidation products was also performed via ultra-high-performance liquid-chromatography (UHPLC Nexera, Shimadzu, Kyoto, Japan) coupled to an electrospray ionization time-of-flight mass spectrometry (ESI-TOF-MS, Triple-TOF^®^ 6600, Sciex, Concord, USA). Samples were separated on a C18 column (Acquity BEH 100 x 2.1; 1.7 µm, Waters) using a mobile phase composed of (A) water with 0.1% formic acid and (B) acetonitrile with 0.1% formic acid. The following solvent gradient was used for sterol oxidation analysis: 50 to 100 % B in 4.5 min, 100 % from 4.5-8.5 min, 100 to 50 % B from 8.5-9.0 min, and 50% B from 9-13 min. MS data were acquired in positive ionization mode. All data were obtained in data-dependent acquisition mode (IDA^®^, Information Dependent analysis) using Analyst^®^ 1.7.1 software.

Sterols and their photo-oxidation products were diluted in isopropanol and centrifuged at 1500 rpm for 5 minutes before injection. The following precursor ions were selected for MS/MS analysis: m/z 411 (ergosterol hydroperoxide, EHP), m/z 429 (5α,8α-epidioxy-22E-ergosta-6,9(11),22-trien-3β-ol, EEP), m/z 377 ((22E)ergosta-5,7,9,22-tetraen-3β-ol , DHE) and m/z 380 (Erg) were monitored, and the data were processed using the Peak View software.

#### Identification of oxidation products by 1H-NMR

The ^1^H NMR (500.13 MHz) spectra were recorded at 0 or 25 °C, using CDCl_3_ as solvent. The analyses were performed on a Bruker-Biospin Ascend 500 instrument, Avance III series, (Rheinstetten, Germany), operating at 11.7 T. The instrument was equipped with 5-mm trinuclear, inverse detection probe with z-gradient (TXI) or 5mm TCI CryoProbe. The temperature was controlled by a BCU I accessory. All chemical shifts were expressed in ppm relative to TMS. The data were acquired and processed using TOPSPIN 3.5 software (Bruker-Biospin, Rheinstetten, Germany).

#### Stability test of ergosterol and endoperoxides

Sterol stock solutions and endoperoxide solutions were dried with N_2_ and resuspended in chloroform, maintaining a concentration of 1 mM. These vials were placed in a thermomixer for 5h at 37°C under stirring at 600 rpm. The aliquots were collected at times zero, 1h, 2h, 3h, 4h and 5h and analysis were carried out by HPLC-DAD.

### Analysis of liposome membrane leakage using a carboxyfluorescein system

#### Liposome membrane leakage with 1,9-dimethyl-methylene blue (DMMB)

Liposomes containing phosphatidylcholine and sterols were prepared by a method adapted from Miyamoto *et al.* (2012) (23). Membranes containing 50 mM carboxyfluorescein (CF) were prepared by mixing POPC (1-palmitoyl-2-oleoyl-sn-glycero-3-phosphocholine) and sterols (Erg, cholesterol, and 7-DHC) in 10 mM Tris buffer at pH 8. Four liposomes were prepared: one with 100% POPC as a control and three others combining 75% POPC with 25% of each steroid. Solutions containing each lipid mixture were placed in a PYREX^®^ test tube, vortexed, and dried under N_2_ stream. The samples were then left for approximately one hour in a vacuum desiccator to ensure the complete removal of solvent. Then, the samples were redissolved in 500 µL of Tris buffer (10 mM, pH 8) containing carboxyfluorescein (50 mM). The resulting mixture was extruded with Avanti Mini-Extruder kit (Avanti Polar Lipids, Alabaster, US) equipped with a 100 nm pore membrane to produce large unilamellar vesicles (LUVs) of 100 nm size.

CF-loaded LUVs were subjected to a Sephadex G-50 column purification using 10 mM Tris with 0.3 M NaCl at pH 8 as a running buffer. The CF-loaded fraction was collected in a microtube and then placed in a 96-well plate at a final concentration of approximately 100 µM of each liposome. DMMB (15 µM) was added to each well and irradiated with a 633 nm LED for 90 minutes. CF externalization was monitored over time with Agilent’s Synergy H1 Hybrid Reader equipment, using excitation at λex = 480 nm and emission at λem = 517 nm. At the end of the reaction, Triton X-100 was added to release all the CF content from the liposomes, to generate the final reading intensity (24, 25).

#### Liposome membrane leakage by oxidation using iron/ascorbate

Liposomes containing egg yolk phosphatidylcholine (EggPC) and sterols (Erg, cholesterol, and 7-DHC) were prepared in Tris-HCl buffer as described in the previous section, however, the DMMB solution was replaced by Fe (III)/ascorbic acid (Fe/Asc) at 10 µM/100 µM final concentrations. All wells contained the final concentrations of approximately 100 µM of each liposome. CF release was measured as described above.

### Quantification of generated oxidized products by HPLC and Mass spectroscopy

#### Quantification of oxidation products from liposome membrane leakage test by HPLC

Liposomes were produced following the same protocol described previously without CF. Briefly, 200 µL of each liposome and 27 µL of DMMB 15 mM were placed in a well plate and irradiated at 633 nm for 90 minutes. Then, the samples were analyzed by HPLC with C8 column (Luna Phenomenex, 250 x 10 mm, 5 µm, 100 Å), isocratic mode (94% MeOH and 6% water); flow rate of 1.0 mL/min at 36°C and 30 µl sample injection volume.

#### Quantification of liposome phospholipids by mass spectrometry

POPC was analyzed by LC/MS in the negative ionization mode, with a C18 column (CORTECS Waters 2.1 mm x 10 mm; 1.6 µM) at 35 °C. The flow rate was 0.2 mL/min; injection volume was 2.0 µL, and the mobile phase was (A) water: acetonitrile (60:40); (B) isopropanol: acetonitrile: water (80:10:2) with ammonium acetate (10 mM) in both phases. The gradient started at 40% and increased to 100% over the first 10 minutes. It was held at 100% from 10 to 12 minutes, then decreased back to 40% between 12 and 13 minutes, and was maintained at 54% from 13 to 20 minutes.

Dimyristoyl phosphatidylcholine (DMPC) (PC14:0/14:0) was used as an internal standard to semi-quantify POPC, and its concentration was calculated by the peak area of the compound divided by the peak area of the internal standard with known concentration. Aliquots of the samples (10 µl) were mixed with 20 µL of DMPC (5 mM) and 170 µL of isopropanol.

#### Quantification of ergosterol, 7-dehydrocholesterol, and their oxidized products by mass spectrometry

To quantify Erg, a calibration curve was built using 7-dehydrocholesterol (7-DHC) as an internal standard. Working solutions for the calibration curve were prepared at concentrations of Erg ranging from 15.6 µM to 1000 µM. To quantify Erg oxidation products (EEP, EHP), a calibration curve was prepared with concentrations ranging from 0.08 µM to 20 µM, using oxidized standards prepared in the laboratory. 7-DHC endoperoxide (CEP) and 7-DHC hydroperoxide (CHP) were used as internal standards at final concentrations of 7.5 µM and 5 µM, respectively.

#### Quantification of cholesterol and cholesterol hydroperoxides by iodometry and HPLC

Cholesterol hydroperoxides (ChOOH) synthesized by photo-oxidation with methylene blue were quantified by iodometry (26). The quantification of cholesterol and ChOOH in the liposome after photo-oxidation with DMMB was carried out by HPLC, following methods described in **Supplementary Materials**.

### In vitro A375 cell viability assay

#### Cell cultures

A375 cells were cultured with Dulbecco’s Modified Eagle’s Medium-high glucose (DMEM, catalog number 10569010, Thermo Fischer Scientific) supplemented with 1% penicillin-streptomycin and 10% FBS. Cells were maintained at 37 °C under a water-saturated atmosphere containing 5% CO_2_, until 80%–90% confluence was reached. Cell counting was performed microscopically with Trypan blue exclusion (27).

#### Photodynamic treatment (PDT) with sterols and DMMB

Cells were incubated in a 96-well plate with increasing concentrations of sterol (Erg, 7-DHC, EEP, or CEP) for 22 h. Then, DMMB (0.1 to 5 nM) (Sigma-Aldrich, São Paulo, Brazil) was added and incubated for an additional 2 h. The medium was then replaced with PBS, and the cell plate was irradiated at final dose of 5J/cm^2^, using red light with 27.84 mW/cm^2^ at 660 nm λ_max_ (light source from Biolambda, Brazil).

#### Cell viability assay

The medium of the cell plate incubated after 0 h and 24h irradiation was discarded and 0.01 g/L resazurin solution in the medium was added to each cell well. Cell viability was determined by measuring fluorescence using the program SoftMax Pro 5.4.1, at 540 nm excitation and 590 nm emission wavelengths. The viability curve was prepared using GraphPad Prism 8.0.1. Data was expressed as means ± standard deviation of at least three independent experiments.

## RESULTS

### Photooxidation of ergosterol and 7-dehydrocholesterol yields endoperoxides as the most stable products

Erg and 7-DHC were photo-oxidized in chloroform with methylene blue as the photosensitizer (**Figure S1**). Aliquots of this reaction mixture were analyzed by TLC and HPLC. Nearly all the sterols were consumed after 90 minutes of irradiation, resulting in the formation of 3 major photo-oxidized products. Oxidized products were first purified by flash column chromatography and fractions checked by TLC (**Figure S2**). The fractions containing the endoperoxide, hydroperoxide, and triene products were dried and resuspended in 3 mL of isopropanol for further HPLCpurification. **Figure S3** shows the peaks corresponding to the three major products that were purified by semi-preparative HPLC and their estimated yields. The photooxidation of ergosterol and 7-dehydrocholesterol primarily produced endoperoxides (EEP and CEP) with yields of approximately 37–38%. Comparatively, the formation of hydroperoxides (EHP and CHP) and triene products resulted in lower yields.

After purification, each oxidation product appeared as single peaks in the HPLC chromatogram, confirming their successful purification (**Figure 2**). Their identities were further confirmed by high-resolution LC-MS analysis.

**Figure 2.**
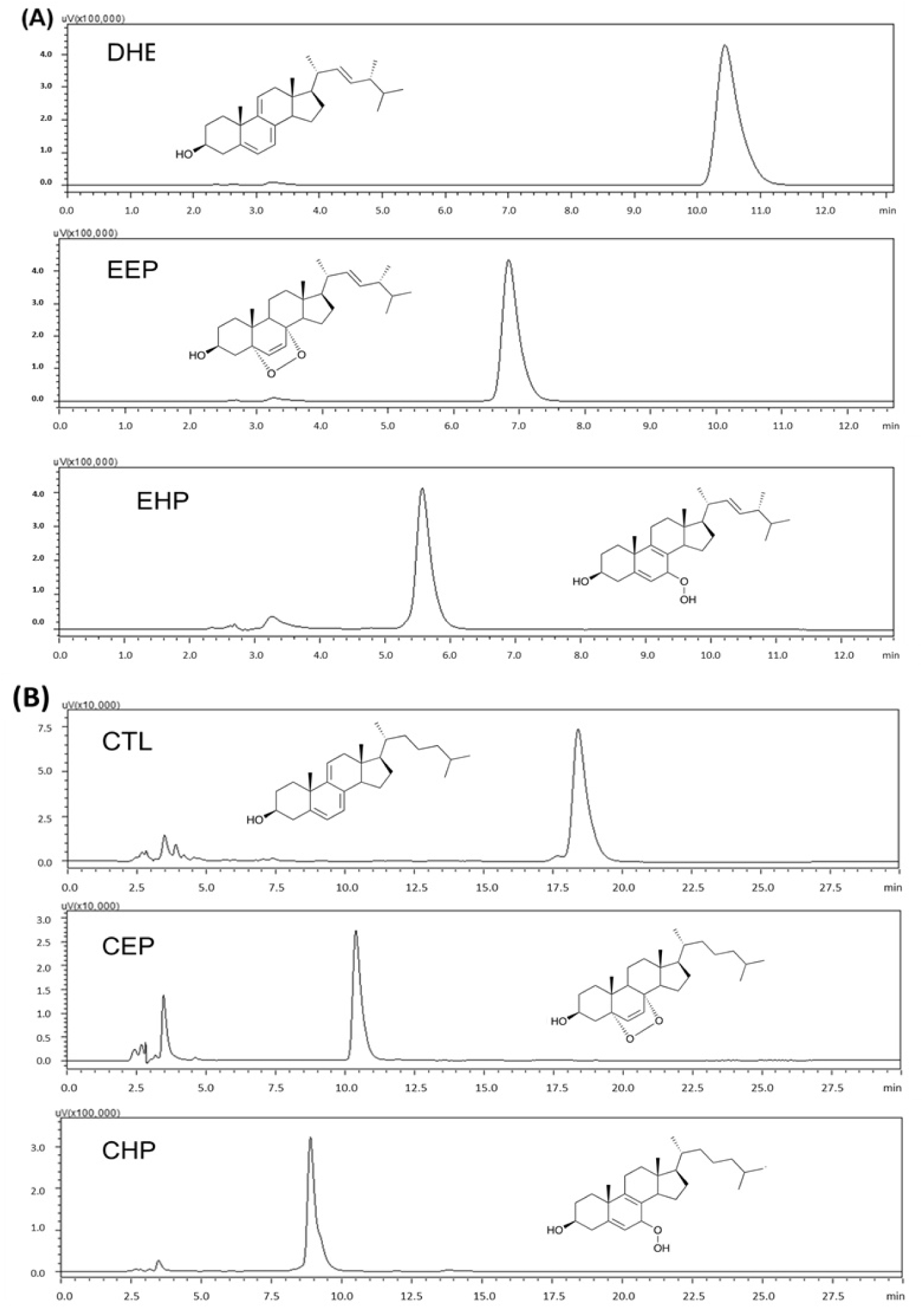
HPLC analysis of the purified photooxidation products derived from (A) ergosterol: dehydroergosterol (DHE), ergosterol endoperoxide (EEP), and ergosterol hydroperoxide (EHP); (B) 7-dehydrocholesterol: cholestanetrienol (CTL), 7-dehydrocholesterol endoperoxide (CEP), and 8-dehydrocholesterol hydroperoxide (CHP).

MS/MS spectra displaying the individual fragmentations of the purified ergosterol oxidation products are shown in **Figure 3**. EEP and DHE protonated ions [M+H]^+^ were observed at m/z 429.3372, respectively, while for the DHE and EHP a dehydrated protonated ions [(M-H_2_O)+H]^+^ appeared as a major ion at m/z 377.3157 and 411.3257. Characteristic fragments were found at m/z 411.32, 393.31 corresponding to the loss of water and O_2_ from EEP; and m/z 375.30 for EHP, corresponding to the loss of three water molecules. Endoperoxides of ergosterol and 7-DHC were further confirmed by 1H-NMR analysis (**Figure S4**, **Table S1** and **Table S2**).

**Figure 3.**
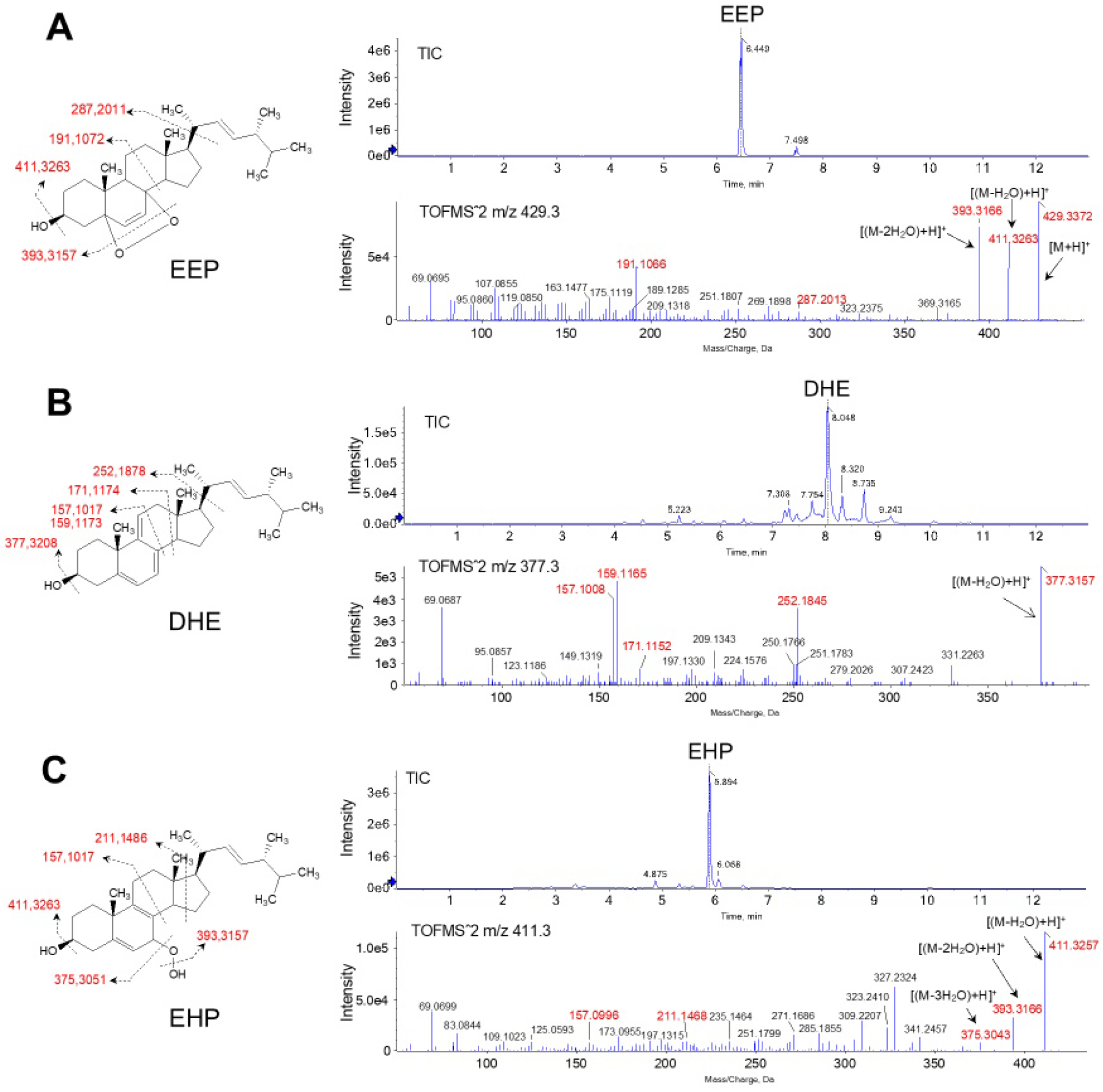
Extracted ion chromatogram and fragmentation spectra obtained in MS/MS mode of Ergosterol photooxidation products: (A) EEP; (B) DHE and (C) EHP.

Next, we investigated the production of endoperoxides and hydroperoxides using dimethyl-naphthalene endoperoxide (DMNO2) as a clean source of ^1^O_2_. Analysis of the oxidation products by HPLC showed that ergosterol incubation with DMNO2 generates both EEP and EHP, suggesting that ^1^O_2_ reacts with ergosterol double bonds through *ene* and Diels-Alder Cycloaddition reactions (**Figure 4**).

**Figure 4.**
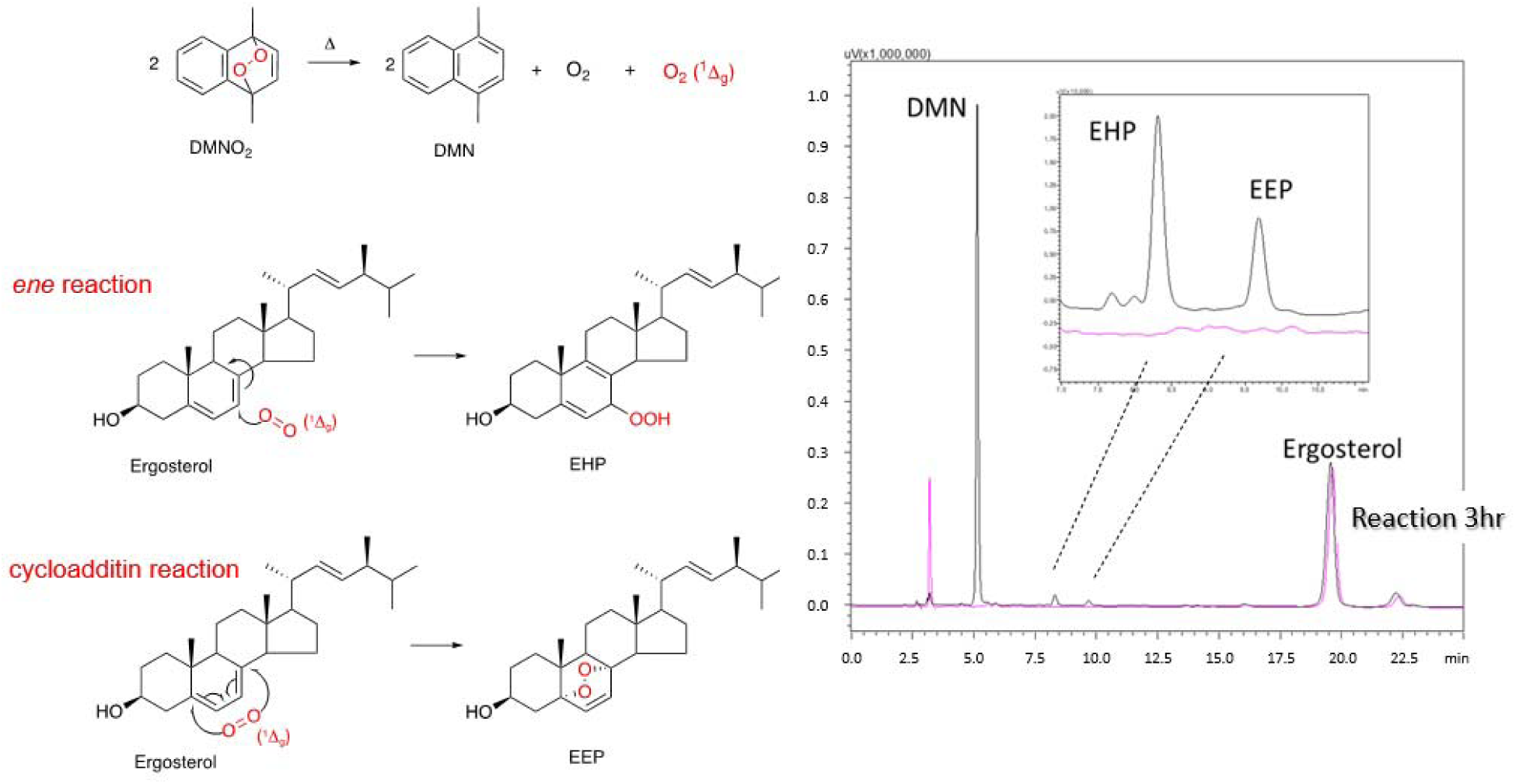
Chromatogram of HPLC analysis of ergosterol reaction with DMNO_2_ monitored at 210 nm. The pink line represents the initial time and the black line the final time of the reaction.

Quantitative analysis of ergosterol oxidation products with increasing concentrations of DMNO2 was performed by mass spectrometry (**Figure 5**). As expected, ergosterol consumption and EEP formation increased proportionally with the generation of ^1^O_2_. However, higher levels of 1O_2_ generated by DMNO2 led to a decrease in EHP levels and an increase in the formation of a third product, identified as a di-hydroperoxide (di-EHP) (**Figure S5**). These results demonstrate that the reaction of ^1^O_2_ with ergosterol produces EEP, EHP, and small amounts of di-hydroperoxides (**Figure 5**). Furthermore, stability analysis showed that EEP is hig1O2hly stable, while EHP degrades rapidly (**Figure S6**). This suggests that EHP is formed through Type II photooxidation but is either converted to di-EHP or degraded during reaction. Overall, these findings indicate that endoperoxides are the most stable products of Erg photooxidation.

**Figure 5.**
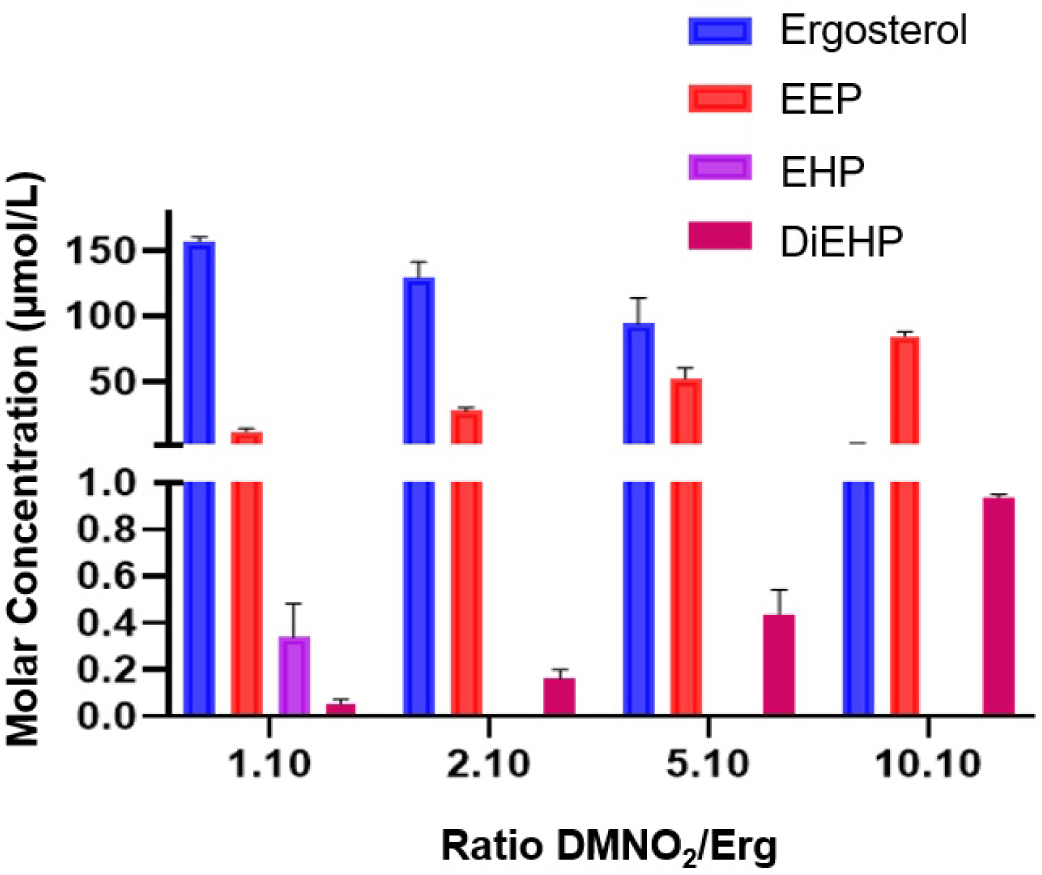
Concentration of ergosterol and its oxidation products analyzed by HPLC-MS after reaction with DMNO_2_.

### Sterols in liposomes exhibit distinct membrane leakage kinetics under type I and II oxidation

Membrane protective effects of Erg and 7-DHC were evaluated under type I and type II oxidation conditions using sterol-containing liposomes loaded with carboxyfluorescein. For type II oxidation (^1^O_2_), liposomes were prepared with POPC (control) or POPC with 25% sterol (Erg, 7-DHC, or cholesterol) and DMMB as photosensitizer. A significant reduction in membrane leakage was observed for liposomes containing sterols (**Figure 6A**). While carboxyfluorescein release in control liposomes reached 75% after 180 minutes of irradiation, liposomes containing Erg or 7-DHC released less than 35% of the probe. Notably, cholesterol provided the highest protection, with minimal release of carboxyfluorescein (<5%). For comparison, membrane leakage studies were conducted using iron/ascorbate (Fe/Asc) as a radical generator under type I oxidation. In type I, lipids are oxidized by radical-mediated reaction, leading to the sequence of initiation, propagation and termination phases. Under the presence of iron and ascorbate as reductant, lipid hydroperoxides (ROOH) react with Fe^3+^ and generate peroxyl radicals (ROO• + Fe^2+^). Subsequently, ROOH react with Fe^2+^ and generate alkoxyl radicals (RO• + Fe^3+^), triggering a cycle of propagation phase of chain reaction (28). Here, in contrast to the results observed under type II oxidation, Erg and 7-DHC provided greater protection to liposomes than cholesterol, particularly during the initial phase of oxidation propagation (**Figure 6B**), which is consistent with their high radical-trapping activity.

**Figure 6.**
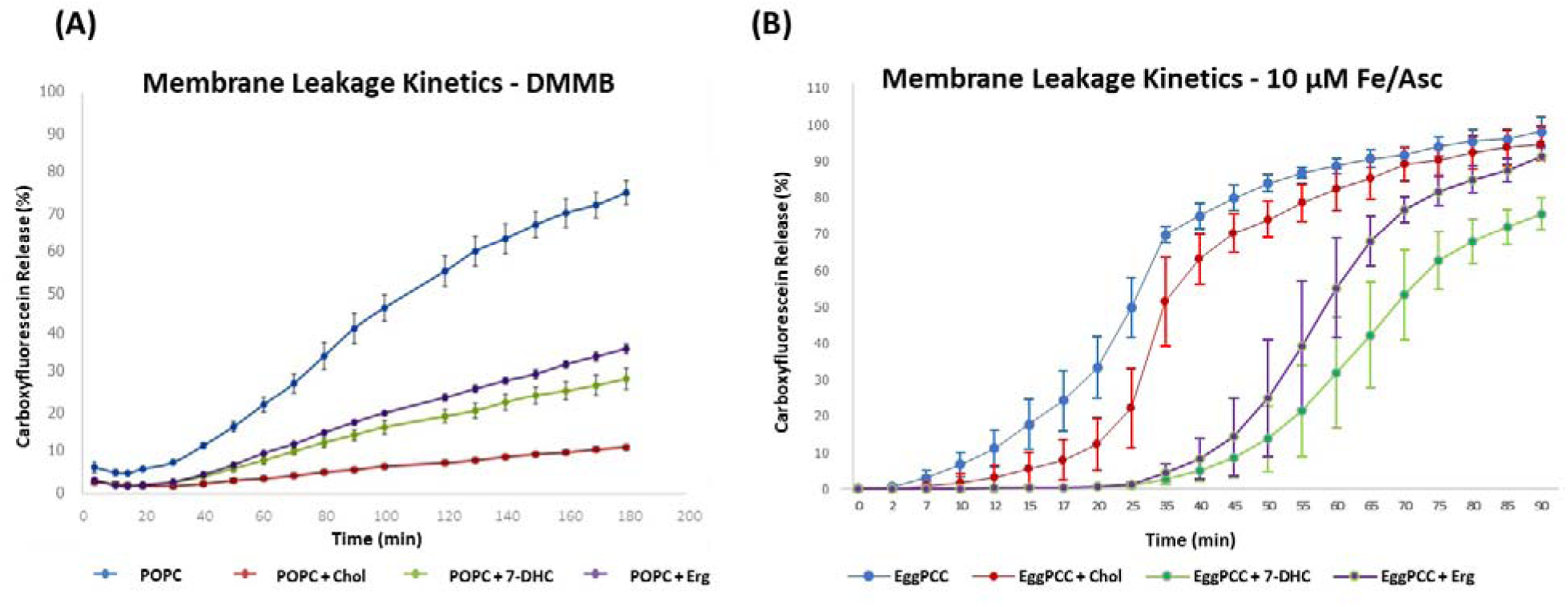
Leakage kinetics of carboxyfluorescein encapsulated in (A) POPC liposome in the presence of the photosensitizer DMMB and (B) Egg phosphatidylcholine liposome (EggPC) in the presence of 10 µM Fe/Asc. Liposomes were composed of either 100% POPC or EggPC, or a mixture of 75% POPC or EggPC with 25% sterols (Erg, 7-DHC or Cholesterol). Leakage was monitored by measuring fluorescence at 517 nm.

### Quantification of phospholipid and sterol photo-oxidation products

#### Sterols protect liposome membranes against phospholipid oxidation

To investigate the membrane protective effects of Erg against photosensitized oxidation, we analyzed the kinetics of phospholipid oxidation in liposome membranes. Phospholipid hydroperoxides (POPC-OOH), the primary products formed by the reaction of ^1^O_2_ with POPC, were quantified by HPLC. As shown, the control decrease (**Figures 7A)** was accompanied by an increase in POPC-OOH (**Figures 7B)**. In control liposomes (POPC), hydroperoxide levels reached a plateau at 20 minutes and then decreased after 40 minutes of irradiation (**Figure 7B**), likely due to the degradation of POPC-OOH forming truncated phospholipid species responsible for membrane leakage (19). In contrast, Erg- and cholesterol-containing POPC liposomes were partially protected from phospholipid oxidation, leading to lower POPC-OOH levels compared to control POPC liposomes. Interestingly, Erg-containing POPC liposomes showed the least generation of POPC-OOH (**Figure 7B**).

**Figure 7.**
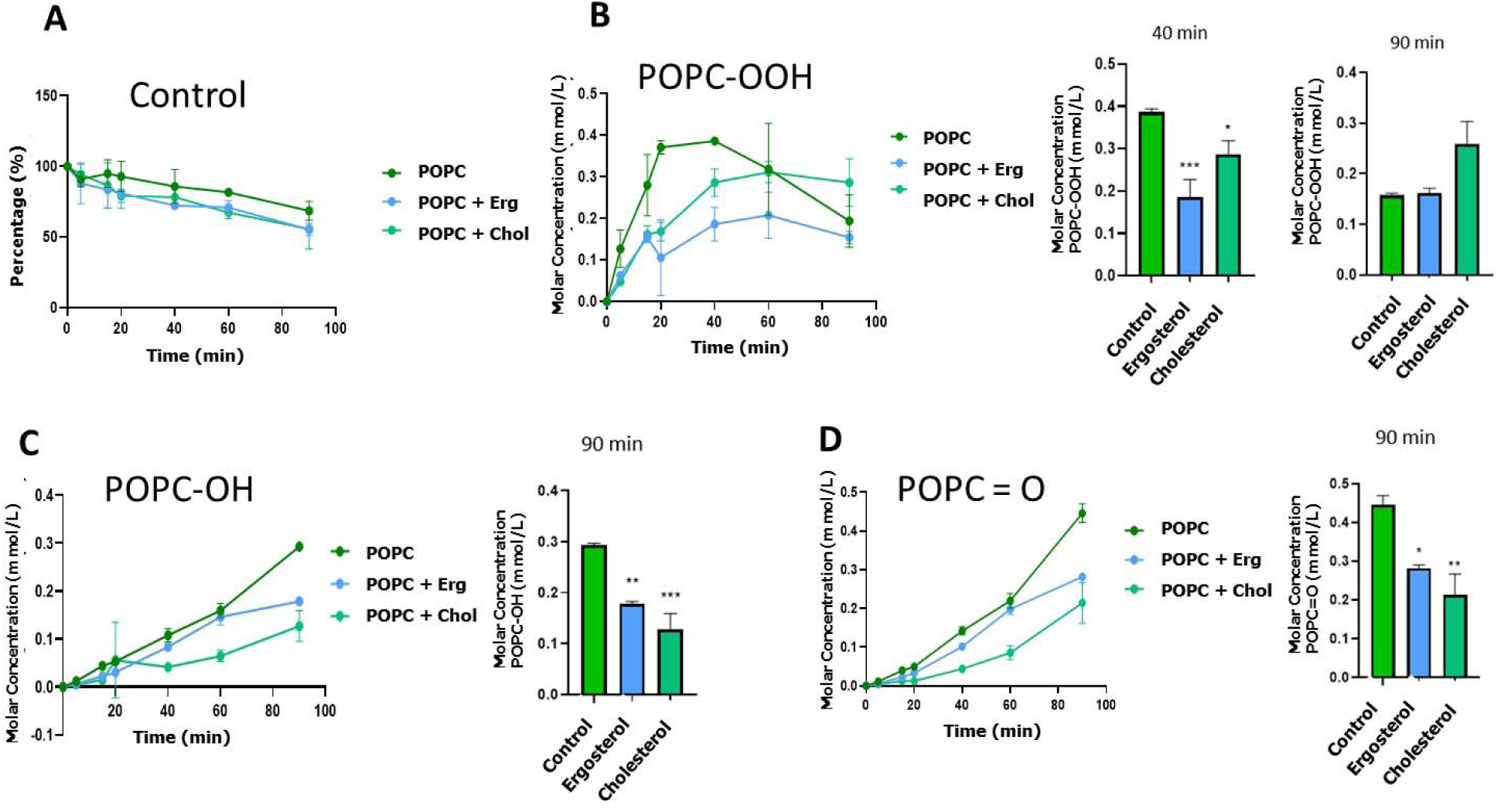
Time-dependent analysis of POPC consumption and formation of photo-oxidation products in liposomes containing POPC (100%), and POPC with cholesterol (Chol) or ergosterol (Erg) (75:25 %). (A) Percentage of POPC over the different reaction times with control. Generation of POPC oxidized products: (B) hydroperoxide (POPC-OOH), (C) hydroxide (POPC-OH) and (D) ketone (POPC=O)

#### Oxidation product consumption kinetics differs among sterols

To determine whether the membrane protective effects of sterols are due to their action as antioxidants or membrane stabilizers, we quantified the consumption of sterols and the formation of oxidized products. Notably, Erg was almost totally consumed within the first five minutes of the photooxidation reaction. This consumption was paralleled to the formation of EEP and EHP, with EEP as predominant product (**Figure 8A**). This indicates that Erg acts as a sacrificial antioxidant, protecting membranes from photodamage. Additionally, this data confirms that EEP is the most stable Erg photooxidation product.

**Figure 8.**
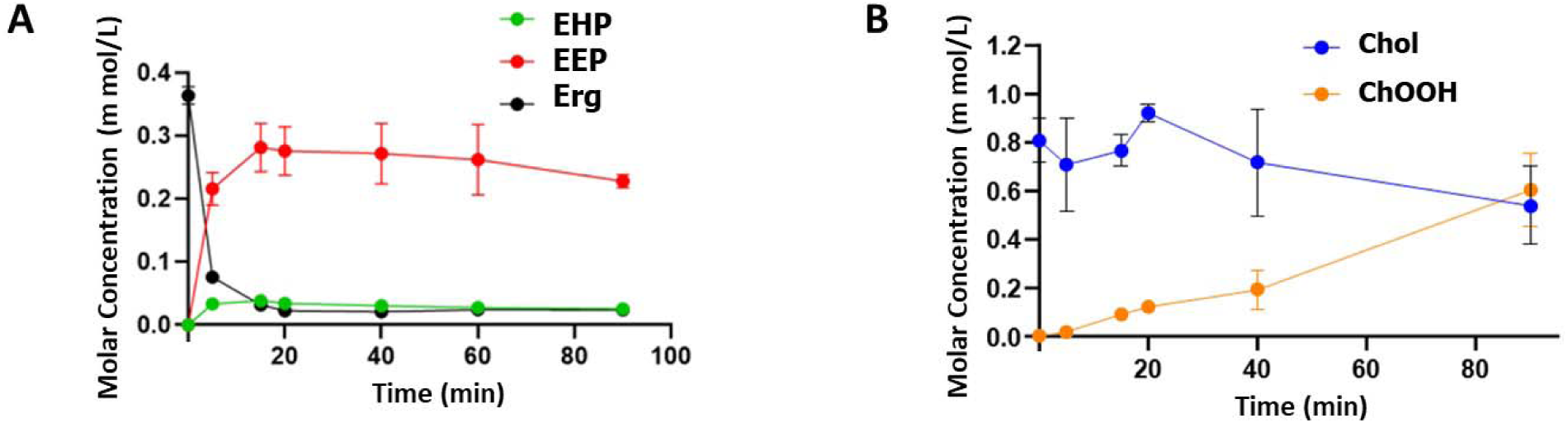
Consumption of sterols and oxidation products formed in the reaction with POPC liposomes in the presence of DMMB. Values obtained by peak integration of the detected products in UPLC-MS using the PeakView program. (A) Erg: ergosterol; EHP: ergosterol hydroperoxide; EEP: ergosterol endoperoxide. (B) Chol: cholesterol; ChOOH: cholesterol hydroperoxides. POPC (100%); POPC with cholesterol (Chol) or ergosterol (Erg) (75:25 %)

In contrast to Erg, the consumption and formation of cholesterol oxidation products showed a different profile. While Erg was consumed within the first few minutes, cholesterol consumption and product formation began only after 20 minutes of reaction. Significant formation of cholesterol hydroperoxides (ChOOH) occurred only after 60 minutes, showing that cholesterol is not rapidly consumed in the reaction (**Figure 8B**). These data suggest that cholesterol protects membranes against photooxidation primarily through its membrane-stabilizing properties, which shield phospholipid acyl chains from ^1^O_2_-mediated oxidation.

### Photo-oxidized products of sterols are superior than their precursor to reduce A375 cell viability

As previously shown, considering EEP the most representative endoperoxide generated by ^1^O_2_ (type II reaction), we compared the A375 cell viability supplementing Erg and 7-DHC and their respective endoperoxides, EEP and CEP. We carried out the assay in four different conditions: (A) PDT, with DMMB (0.1 nM) and irradiation (5 J/cm^2^); (B) with DMMB (0.1 nM) and without irradiation to evaluate the dark toxicity of DMMB; (C) without DMMB and with irradiation (5 J/cm^2^) to evaluate the effect of irradiation; and (D) without DMMB nor irradiation as a control. **Figure 9** shows the significant reduction in cell viability of A375 cells treated with both endoperoxides when compared to their precursor molecules. We did not observe significant effects of dark toxicity (**Figure 9B**) or irradiation alone (**Figure 9C**) on cell viability. This experiment clearly evidenced the negative viability effects of Erg and 7-DHC endoperoxides in A375 cells under PDT as compared to their precursors.

**Figure 9.**
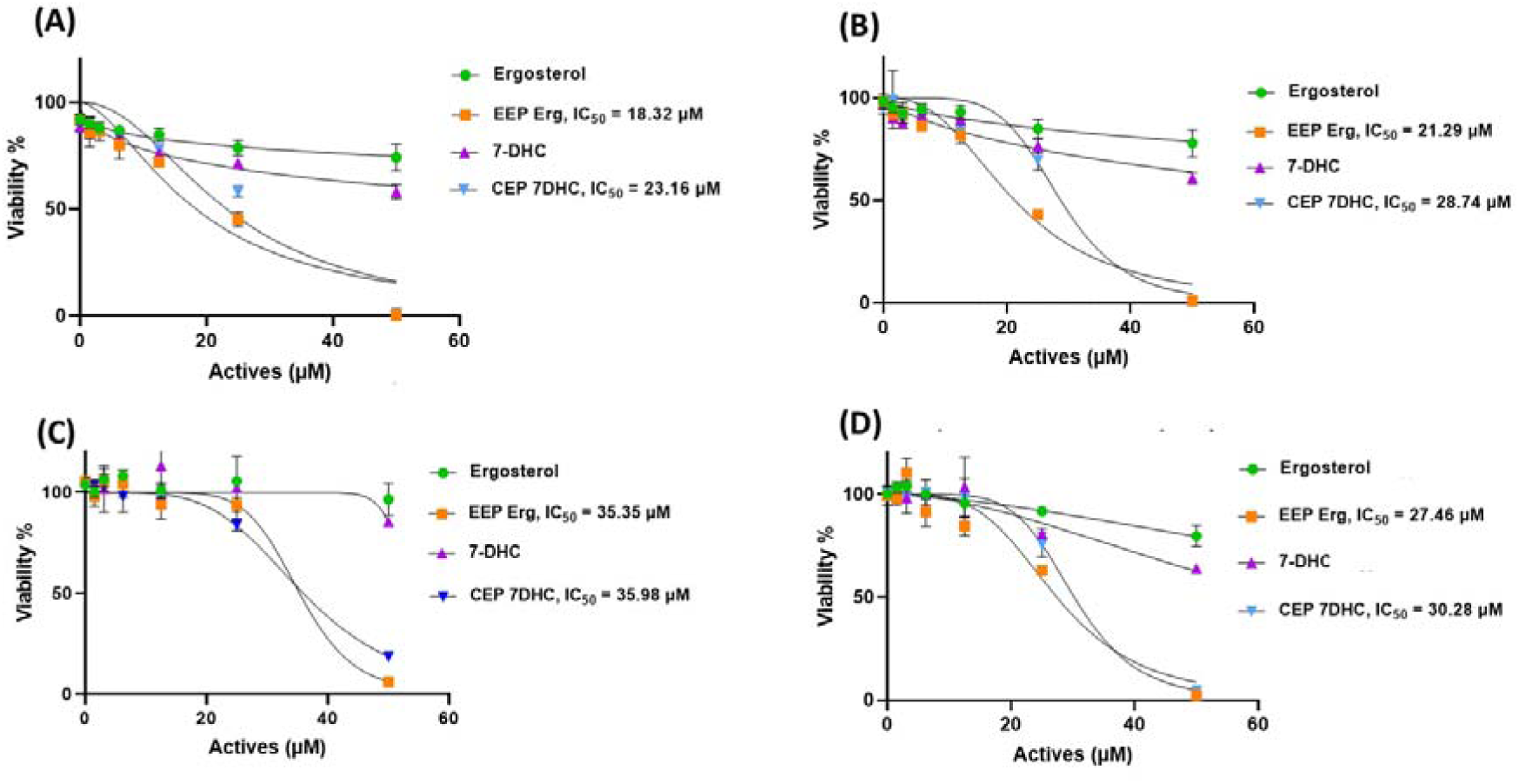
Cell viability effect in A375 with different sterols, ergosterol, ergosterol endoperoxide (EEP Erg), 7-DHC and 7-DHC endoperoxide (CEP): (A) PDT, with DMMB (0.1 nM) and irradiation (5 J/cm^2^); (B) with DMMB (0.1 nM) and without irradiation; (C) without DMMB and with irradiation (5 J/cm^2^); and (D) without DMMB and irradiation.

## DISCUSSION

### Oxidation products of ergosterol by singlet oxygen (^1^O_2_) reaction

Our protocol of type II reaction (^1^O_2_) by DMMB was different than used in other studies. For instance, Böcking *et al*. (2000) (29) used toluene blue as a photosensitizer to generate ^1^O_2_ in *S. cerevisiae* and this resulted in the yield of 4.0% DHE followed by 1.5% EEP. Ponce *et al.* (2002) (30) carried out several reactions with rose Bengal and eosin with three types of solvents (pyridine, ethanol, and methyl *tert*-butyl ether), and EEP was the only product generated in all combinations. In the latter study, Erg + methyl *tert*-butyl ether + eosin generated 16% EEP and, 8% EHP, with trace amounts (not quantifiable) of DHE. According to the characterization data by NMR and mass spectrometry, we confirm that the oxidation products synthesized and purified in our work are the same as those found in the literature, regardless of solvent or photosensitizer types. As mentioned in both our results and in the literature, EEP was the most abundant product in type II Erg oxidation reaction (29–31). Thus, this molecule is a good marker for ^1^O_2_ reaction given its stability at high temperatures (29, 32).

### Membrane leakage evaluation using DMMB as a photosensitizing agent

In this experiment, the purpose was to investigate membrane leakage caused by the photo-oxidation type II reaction (^1^O_2_) on phospholipid and sterol. Cholesterol-containing POPC liposomes showed lower membrane leakage over time compared to control, Erg or 7-DHC POPC liposomes (**Figure 6A**). This result confirmed the different protection efficacy promoted by sterols, corroborating data from Tsubone *et al.* (2019) and Bacellar *et al.* (2014) (33, 34), for instance, showed that a liposome formed by dioleoyl-*sn*-glycero-3-phosphocholine (DOPC) with 30% cholesterol decreases its permeability by up to 60% under photo-oxidative stress. Additionally, a study on molecular dynamics by Zhang *et al.* (2018) (35) reported the absence of pore formation in cholesterol-rich membranes under photo-oxidative stress. These authors suggested that cholesterol possibly formed hydrogen bonds with the oxidized lipids, inhibiting membrane permeability loss. While membrane protection of POPC liposomes containing Erg and 7-DHC was observed, the effects are clearly inferior than those containing cholesterol. A likely explanation is the rapid oxidation of Erg and 7-DHC by type II reaction, leading to reduced membrane stabilization effect as compared to cholesterol.

After some time of reaction, however, all liposomes began to display leakage possibly due to formation of truncated phospholipid species bearing aldehyde and carboxylic acid groups from phospholipid hydroperoxides (**Figure 7B**). These truncated lipids favor the deformation of the bilayer and thereby the formation of pores, increasing membrane permeabilization (33).

### Membrane leakage evaluation using Fe/Asc as oxidizing agent

For comparison, iron/ascorbate (Fe/Asc) was used as a membrane oxidant to verify the protective efficacy promoted by sterols in liposomes against damage induced by free radicals (type I reaction). Asc reduces Fe (III) into Fe (II), and the formation of Fe (II) initiates a Fenton-type reaction with peroxides, which produces alkoxyl radicals (27). Using 10 µM Fe/Asc, a greater membrane protection using Erg and 7-DHC was obtained relative to control and cholesterol at the initial phase of lipid peroxidation (**Figure 6B**). Nonetheless, at the propagation phase, this difference among sterols was no longer significant.

A study carried out by Freitas *et al.* (2024) (22) also noted that 7-DHC protects the membrane, as it reacts to produce two non-enzymatic 7-DHC oxidation products. This indicates a preferential oxidation of 7-DHC relative to phospholipids, preventing the formation of truncated species. Although an in-depth study for Erg has not been conducted yet, the similarity of this molecule to 7-DHC (the presence of the conjugated double-bound present in the sterol B-ring) suggests that Erg may have the same membrane protection properties as 7-DHC. Indeed, these authors validated this hypothesis in yeast strains with targeted deficiencies of genes important for Erg biosynthesis.

### Oxidation products of liposomes generated by DMMB photooxidation

Analysis of membrane oxidation products showed that concentrations of phospholipid hydroperoxides (POPC-OOH) were higher in control membranes than in Erg- or cholesterol-containing POPC liposomes (**Figure 7B**). While the formation of hydroperoxides, hydroxides, and ketones in the acyl chain of phospholipids can alter the area occupied per lipid in the bilayer, these modifications likely do not lead to membrane disruption. Previous studies conducted by Bacellar et al 2019 (36) and Freitas et al 2024 (22) suggested that membrane disruption is caused by truncated phospholipids containing fragmented acyl chains, such as POPC aldehyde (AldoPC) or POPC carboxylic acid (PAze PC). These molecules have a different conformation as compared to phospholipids with two intact rather than fragmented acyl chains. In fact, truncated phospholipids present a shorter chain, leading to a more conical conformation, which may facilitate membrane leakage (36).

Thus, the ^1^O_2_ attack to membrane phospholipids generates hydroperoxides by the ene-reaction. Since the hydroperoxide groups in the phospholipid acyl chain tend to migrate to the polar head region of the membrane bilayer, they change and decrease the membrane thickness, but this is not enough to promote membrane disruption (36). Once formed, phospholipid hydroperoxides can undergo one-electron reduction and oxidation reactions, either through Type I mechanisms or in the presence of metal ions, generating alkoxyl and peroxyl radicals (6, 22, 37). These radicals can initiate membrane chain peroxidation reactions that increases hydroperoxide formation. Additionally, alkoxyl radicals β-scission, induces phospholipid acyl chain cleavage generating truncated phospholipids bearing aldehyde and carboxylic acids, which have strong effects on membrane bilayer permeability due to their molecular conformation, hydrogen bonding capabilities and chain mobility (36, 38, 39), ultimately resulting in a much thinner membrane, decreased membrane packing and formation of transmembrane pores (37).

Here, the formation of truncated phospholipids, such as AldoPC was not apparent in liposomes analyzed by UPLC-MS analysis. However, in the control sample (POPC), the maximum level of POPC-OOH were observed between 20 and 40 minutes, followed by a decline, suggesting that POPC-OOH was consumed after this reaction time (**Figure 7B, POPC**). The consumption of hydroperoxides in oxidation reactions is an indication of radical reactions (type I oxidation), caused by the photosensitizer, which can generate truncated lipids such as AldoPC and PAze PC (28). Although POPC liposomes containing cholesterol (**Figure 7B, POPC + Chol**) produced more POPC-OOH than those with ergosterol (**Figure 7B, POPC + Erg**), there was no indication of ChOOH or POPC-OOH reduction over the time (**Figure 8B**). The post-irradiation one-electron reduction of ChOOH, similar to POPC-OOHs, can induce the formation of radicals capable to initiate and propagate chain peroxidation reaction. However, membrane leakage kinetics tests under ^1^O_2_ exposure showed that POPC + Chol liposomes had the lowest membrane leakage among the samples tested (**Figure 6A**). This data indicates that, under our experimental conditions, hydroperoxides in membranes containing cholesterol was shielded from one-electron reactions to a degree that would cause membrane disruption.

This data contrasts from POPC + Erg, which showed a slight decrease in POPC-OOH after 60 minutes of reaction (**Figure 7B, POPC + Erg**), suggesting the formation of truncated lipids. Thus, this difference in the consumption of hydroperoxides, and the generation of truncated phospholipids, may explain why Erg-containing liposomes experience faster membrane disruption than those containing cholesterol (**Figure 6A**). We hypothesize that Erg or 7-DHC initially acts as sacrificial antioxidants, likely with the Fe/Asc free radical oxidation system, by reacting more rapidly with ^1^O_2_ than POPC producing endoperoxides, EEP and CEP. The estimated rate constants for the ^1^O_2_ quenching are 1.2 x 10^7^ M^-1^s^-1^ (40) for ergosterol and 10^4^-10^5^ M^-1^s^-1^ for phosphatidylcholine (11). However, once Erg or 7-DHC is oxidized membranes containing endoperoxides (EEP or CEP) is more susceptible to one electron oxidation or reduction reactions than membranes containing cholesterol and its hydroperoxides.

In addition to differences in the reactivity, Tsubone *et al.* (2019) suggested that cholesterol protection is related to the formation of hydrogen bonds with oxidized lipids, making it difficult to change the angle of inclination of the lipids that are interacting, thus maintaining the integrity of the membrane (33). Other work mentioned that cholesterol can reduce the oxygen permeability in the membrane, since there is a greater electron density on both sides of the membrane bilayer when the amount of cholesterol is increased (41, 42). Packing of cholesterol with the phospholipid layer would thereby generate a physical barrier to the oxygen diffusion (41, 42). Our data showed slower consumption of cholesterol and formation of oxidation product (ChOOH) in cholesterol-containing liposomes (**Figures 8B**) than those containing ergosterol (**Figures 8A)**. Interestingly, even with distinct oxidation reactions and liposomes composition, such as 1,2-dipalmitoyl-sn-glycero-3-phosphocholine (DPPC) and 1-palmitoyl-2-oleoyl-sn-glycero-3-phospho-(1’-rac-glycerol) (POPG) in different studies (33, 35), the same tendency is observed, thus supporting the consistency of our results.

As a complement, an interesting work using plant extract also evidenced the antioxidant and physical adsorption mechanisms of several plants to protect membrane, measured via CF emission in DMMB-liposomes. Gallic acid and trehalose were used as standard compounds, and the percentage of membrane protection was linearly proportional to the concentration of these compounds, highlighting the multiple mechanisms that may influence membrane integrity (43).

### Cell viability effect of sterols and photodynamic therapy in the melanoma A375 cell line

Our results showed significant reduction in cell viability of melanoma A375 cells treated with sterol endoperoxides. EEP and CEP, the endoperoxides of Erg and 7-DHC, respectively, showed greater efficacy in killing melanoma cells compared to their precursors. Higher cytotoxicity of sterol endoperoxides was also reported in other cancer cells. On A549 lung cancer cells, authors showed greater cell viability inhibition of EEP to Erg and 9,11-dehydroergosterol endoperoxide. According to their work, EEP induced caspase-dependent apoptosis via mitochondrial damage, cytochrome c release, and ROS generation (44). Moreover, in the triple-negative breast cancer cellular model, EEP and its derivative (made to improve the aqueous solubility of EEP) demonstrated effectiveness to induce cell death and selectivity without affecting non-cancerous cells (45). Furthermore, the superior cytotoxicity of endoperoxides derived from ergosterol and 7-DHC has been evidenced in lung cancer A547, cervical carcinoma cells HeLa, breast cancer cells SKOV-3, prostatic carcinoma cells DU145, and human normal liver cells L-02 (46).

Interesting studies on 7-DHC by Freitas et al. (2024) have demonstrated its influence in ferroptosis, an iron-dependent and non-apoptotic cell death induced by accumulation of membrane-destabilizing truncated phospholipid species. 7-DHC has been deemed a suppressor of ferroptosis, due to its high reactivity towards peroxyl radicals, which protects polyunsaturated fatty acids (PUFAs) from peroxidation. Although the unsaturated B rings in 7-DHC are highly susceptible to peroxidation, acting as a sacrificial antioxidant, the oxidized 7-DHC metabolites have a lower membrane-destabilizing effect compared to truncated species. This may provide a membrane-shielding effect that reduces cell death. Membrane leakage tests showed superior membrane protection of 7-DHC compared to control and cholesterol against radicals produced by the Fe/Asc system (22). This result is in line with our results of 7-DHC-membrane leakage using Fe/Asc (type I reaction) and carboxyfluorescein.

Another work by Li et al. (2024) detected the location of 7-DHC inside the cells, which was both in the mitochondria and the plasma membrane. These authors demonstrated the 7-DHC as suppressor of phospholipid peroxidation to reduce ferroptosis in both locations. They have also showed that Erg, the analogue of 7-DHC, also suppressed ferroptosis. By adding RSL3 (ferroptosis-inducing agent) to HEK293T cells pre-treated with 7-DHC, Erg, stigmasterol, or cholesterol, they showed that Erg and 7-DHC blocked phospholipid radical-derived peroxidation and inhibited ferroptosis, unlike other sterols (21).

As the mechanism of ferroptosis relies on the high susceptibility of PUFAs from phospholipid membranes to lipid peroxidation, it leads to the accumulation of lipid peroxides and cell death. Nevertheless, it is known that lipid droplets (LDs), the lipid-rich cellular organelles, incorporate excess PUFAs within their neutral core, controlling PUFA storage in the form of triacylglycerols and sterol esters to reduce potential lipotoxic damage (47). It was shown that the formation of LDs induced by cell cycle arrest promoted resistance to ferroptosis. The latter study showed that diacylglycerol acyltransferase (DGAT)-dependent LDs were formed under induction by cell cycle arrest. This LD sequestered PUFAs accumulated in cells to suppress ferroptosis, and the inhibition of DGAT resensitized the cell arrest to ferroptosis (48). Moreover, as LDs disruption may release excessive lipids to generate increased lipid peroxidation, they can act in either prevention or induction of lipotoxicity and cell death, depending on the oxidative stress cell status. There exist data indicating that the ^1^O_2_ generated by PDT can induce both apoptosis and ferroptosis, according to the ^1^O_2_ dose. These authors described that a moderate dose of ^1^O_2_ induces apoptosis, releasing cyt c from mitochondria. However, an excessive dose of ^1^O_2_ inactivates cyt c and caspase enzymes, inhibiting this pathway and inducing ferroptosis by excess lipid peroxidation combined with iron release from lysosomal membrane disruption (49).

Interestingly, it is reported that Erg can also be stored in LD as sterol ester, and this would prevent lipid peroxidation that leads to ferroptosis (50). This study demonstrated that Erg in *S. cerevisiae* is mainly located in the inner mitochondrial membrane (IMM) rather than in the outer mitochondrial membrane (OMM), and that Erg is actively transported to the OMM in an energy-independent way. Since mitochondria closely communicate with LDs, endoplasmic reticulum (ER), and peroxisomes (51), the LD storage of Erg might be influenced by the correct function of mitochondria. In contrast, the endoperoxide EEP has been shown to inhibit LD synthesis of differentiated 3T3-L1 cells (52), indicating the opposite effects of Erg and EEP on LD formation.

Concerning the role of mitochondria, it is well known that these organelles and lysosomes are the main targets for type II reaction (^1^O_2_) by PDT (35). Although lipid peroxidation in the ER and plasma membrane highly influences ferroptosis, mitochondria have also a contribution to this cell death process (53). It is described that PDT with DMMB reduced mitochondrial membrane potential and thereby promoted an increase in reactive oxygen species, autophagic dysfunction, and membrane permeability (51). DMMB easily diffuses through the plasma membrane and, due to its positive charge, accumulates in the mitochondria, an organelle with an inner negative charge (54). It is reported that PDT with DMMB may lead to loss of membrane potential of mitochondria in the parasite *Leishmania amazonensis*, and released lipids into the cytosol (ref). Additionally, LDs increased substantially and accumulated large amounts of released lipids. This response of LDs to mitochondrial dysfunction regulates lipotoxicity and preserves organelle function to prevent cell death. However, a redox imbalance caused by lipid overload possibly led to constant cellular stress, and the parasite’s final death (55).

As a complement, a clinical test with PDT by intratumorally injection of 2% methylene blue aqueous solution with 2% lidocaine, in a 92-years-old multiple skin melanoma patient, showed a tumor remission in five out of six sites, highlighting the potential of PDT as an effective and inexpensive treatment for melanoma (56).

All above studies could provide several insights into the cell death mechanism in A375 cells in this work, as follows:

First, changes in membrane permeability property according to the type of oxidative damage. In the type I reaction, superior membrane protection against leakage was observed for Erg and 7-DHC than for control and cholesterol in the initiation phase of lipid peroxidation. This agrees with the results presented by Freitas *et al.* (2024) (22), indicating the effectiveness of 5,7-unsaturated sterols against type I reaction in the initial phase of lipid peroxidation. This is possibly due to the radical trapping ability (57) and reduced formation of truncated lipids in the membranes where these sterols are embedded (22). This result, however, contrasts with type II oxidation, where cholesterol provided a better membrane protection than Erg and 7-DHC. Interestingly, Erg is almost immediately consumed and generates EEP or EHP, while cholesterol consumption and generation of its main hydroperoxide (ChOOH) takes much longer under type II photo-oxidation. As mentioned previously, cholesterol-rich membranes reported the absence of pore formation possibly due to the hydrogen bonds formation with the oxidized lipids (35).

Second, the protection of cells against ferroptosis in Li *et al.* (2024) and Freitas *et al.* (2024) (21, 22) had a greater effect with 7-DHC than controls. Our results also showed a non-significant cell viability reduction in A375 cells treated with 7-DHC and Erg. Here we can raise some hypotheses. The non-excessive dose of ^1^O_2_ by PDT (type II reaction) in our work led to the formation of a small amount of their respective endoperoxides, not enough to kill the cells, but enough to protect the membranes from truncated lipid formation. Another hypothesis might be the high amount of both sterols that induced the formation of LDs, thereby avoiding lipotoxicity and cell death. Nonetheless, the above hypotheses are yet to be confirmed.

Third, EEP and CEP promoted opposite effects as compared to their precursor molecules, Erg and 7-DHC. These endoperoxides led to cell death, while Erg and 7-DHC did not. According to the work by Wu *et al.* (2018) (44) mentioned previously, EEP induces caspase-dependent apoptosis via mitochondrial damage, cytochrome c release, and ROS generation, and induces mitophagy at 20 µM. Interestingly, our work showed a sudden decline of cell viability mostly after 20 µM EEP (**Figure 9**), which highlights the need for additional studies to provide a deeper understanding of this dose-dependent mechanism. Our results are also be in line with Fujii *et al.* (2023) (49), who suggest that a moderate dose of ^1^O_2_ mainly induces apoptosis rather than ferroptosis. An additional explanation for the reduction of cell viability may be the lipotoxicity due to the excess concentration of EEP. This endoperoxide may suppress lipid droplet synthesis by inhibiting triglyceride synthesis in differentiated 3T3-L1 cells (52). Another study with 24(S)-hydroxycholesterol showed that its esterification resulted in the formation of atypical lipid droplets due to the endoplasmic reticulum membrane disorder (56). This was caused by the change of polarity of this sterol by esterification, disturbing membrane bilayer orientation (58). Although they differ in structure, changes in molecular polarity induced by enzymatic or non-enzymatic processes may lead to non-controlled lipotoxicity and increased cell death. This information may inspire future investigations concerning the influence of oxidation products on the rupture of LDs, thereby promoting lipotoxicity (55). Moreover, EEP and CEP under PDT may generate secondary metabolites. However, as shown in **Figure S6**, EEP is highly stable, suggesting that the effect of PDT on the already-formed EEP may not be highly significant. All these descriptions strongly highlight that these endoperoxides have different efficacy compared to their precursor molecules, and dose and stress conditions may be one of the key triggers to direct cell death pathways.

As a limitation and challenge of our work, we can mention as follows: (1) this study focused on type II photooxidation (^1^O_2_ reaction), therefore, studies of subsequent reactions triggered by chain reactions, with the intervention of type I reaction (radical reaction) in these sterols, remains to be elucidated; (2) it lacks in-depth characterization of hydroperoxides generated by the type II reaction. For example, our study was able to purify the hydroperoxide product formed by ^1^O_2_ attack at C7 and H-abstraction from C9, producing 7-OOH with unsaturated bonds at C5(6) and C8(9). Unfortunately, results for other hydroperoxides were inconclusive, as we could not isolate them in sufficient quantities for NMR analysis and they were also rather unstable; (3) further study is needed to allow extrapolation of membrane leakage tests using phospholipids to those tests using cells. Although this membrane leakage method is widely used, different cancer cells may express different properties, which makes it challenging, for example, to compare different tumor cells based on this same method.

Although further studies are still necessary to confirm these discussions, we described the step-by-step process to generate and identify endoperoxide through the type II reaction, and provided new insights to elucidate different mechanisms of cell death promoted by Erg, 7-DHC and their respective endoperoxides, in combination with PDT in A375 melanoma cells. It is also worth mentioning that the results obtained by endoperoxides EEP and CEP has relevant applicability, thus, the tests can be extended to the animal level.

## CONCLUSION

Melanoma is one of the most aggressive oncological diseases that require effective treatment. Erg is a sterol from a natural source that has attracted attention for its several benefits. 7-DHC, an analog of Erg present in the human membrane, has also shown benefits in several diseases, including cancer. However, some ambiguity or uncertainty remains regarding the mechanism of their protective or deleterious effects. Focusing on the mechanistic point of view of the cell membrane lipid bilayer, we used the liposome leakage model to compare the possible causes of membrane rupture in distinct types of lipid oxidation reactions. We then evaluated the cell viability of A375 melanoma treated with 7-DHC and Erg, and their respective endoperoxides generated by the type II oxidative reaction.

The protocol carried out in this work confirmed the generation of Erg endoperoxide and other sterols. The membrane leakage results demonstrated different behaviours between liposomes composed of Erg, 7-DHC or cholesterol under distinct oxidation reactions (type I and II). When correlating our results with the literature, the presence of truncated lipids generated by oxidative stress appears to be the main cause of membrane integrity disruption, compromising cell viability. In vitro viability testing of A375 melanoma cells with PDT showed inhibition of cell viability in endoperoxide-treated cells and no significant reduction in precursor molecules. These results in combination with data from the literature pointed out some directions, such as the influence of these sterols and derivatives combined with ^1^O_2_ on apoptosis, ferroptosis, mitochondria integrity and formation of lipid droplets. The change in cell viability appears to be highly dependent on the dose of oxidative stress and the concentration or type of sterols.

Thus, in this work, unlike other studies, we explored the possible effects of Erg and 7-DHC oxidation derivatives generated mainly by the type II reaction and highlighted the different effects of sterols and their endoperoxide derivatives on the cell viability of the A375 melanoma cell line. As a future perspective, in association with PDT - which is known to be less invasive than conventional cancer treatment methods such as surgery - these endoperoxides may open up great possibilities for developing new formulations, including nanoparticles, as potential alternatives for melanoma treatment.

## ACKNOWLEDGMENTS

This study was supported by Fundação de Amparo à Pesquisa do Estado de São Paulo [FAPESP, CEPID–Redoxoma grant 13/07937-8 to S.M., and M.S.B, and P.D.M.], [FAPESP, nbr. 2023/15361-0 to M.N.Y.], [FAPESP, 23/12767-6 to T.E.O.S.], Conselho Nacional de Desenvolvimento Científico e Tecnológico [CNPq 313926/2021-2 to S.M.], [CNPq n°. 304350/2023-0 to P.D.M.], Coordenação de Aperfeiçoamento de Pessoal de Nível Superior [CAPES, Finance Code 001] , Pro-Reitoria de Pesquisa da Universidade de São Paulo [PRPUSP], NAP Redoxoma [n°. 2011.1.9352.1.8 tp P.D.M] and the John Simon Guggenheim Memorial Foundation to P.D.M.

## SUPPLEMENTARY MATERIALS

See supplementary materials.

