## Supplemental information for "Comparative study of Ergosterol and 7-dehydrocholesterol and their Endoperoxides: Generation, Identification and Impact in Phospholipid Membranes and Melanoma Cells"

#### SUPPLEMENTARY FIGURES, TABLES AND METHODS

### Generation of sterol oxidation products by photo-oxidation

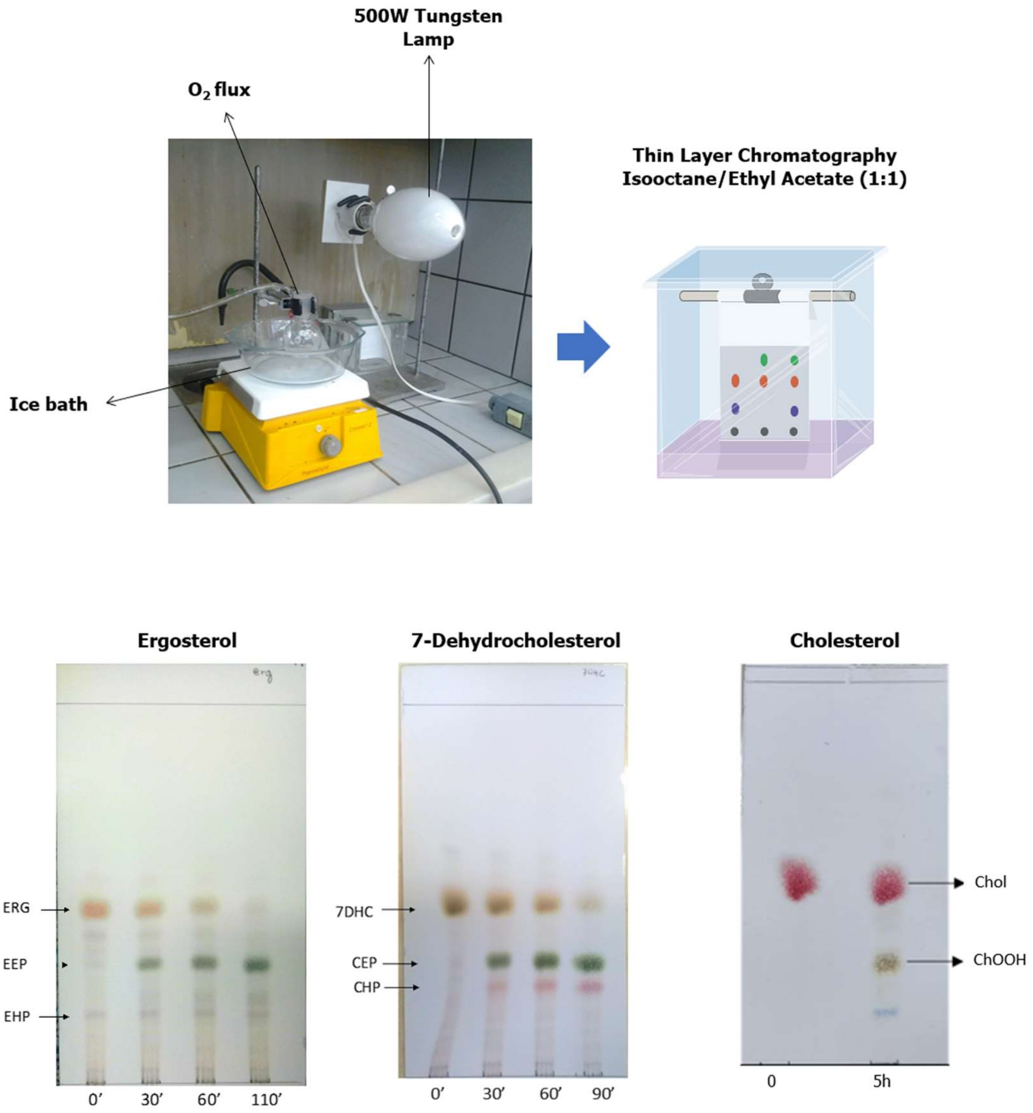


**Figure S1:** Photo-oxidation of sterols and qualitative analysis of their oxidation products by thin layer chromatography (TLC). Sterols were photooxidized in the presence of methylene blue by using a tungsten lamp. Samples were kept cooled by an ice-bath. Aliquots of the reaction mixture were taken and the formation of oxidized products were checked by TLC analysis. Cholesterol (Chol), Ergosterol (Erg), 7-dehydrocholesterol (7-DHC), Ergosterol Endoperoxide (EEP), Ergosterol Hydroperoxide (EHP), Cholesterol Endoperoxide (CEP), 8-Dehydrocholesterol Hydroperoxide (CHP), and Cholesterol Hydroperoxide (ChOOH).

**Purification of sterol oxidation products**

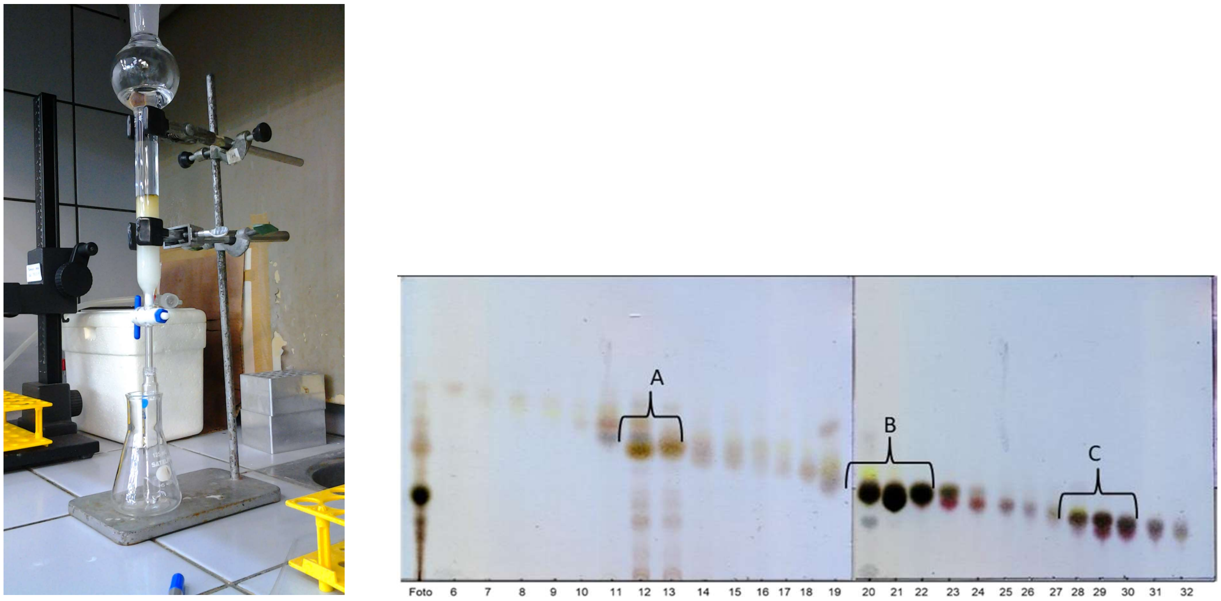


**Figure S2.** Purification of sterol oxidation products by silica gel flash column chromatography and analysis of the fractions by thin layer chromatography. Aliquots of 10 µL of the samples were applied to the plate and eluted with isooctane: ethyl acetate. After elution, the plate was developed using $H_2SO_4$:$H_2O$ (1:1 v/v) spray and heating on a hot plate. The order of elution of the products was (A) Ergosterol Hydroperoxide (EHP), (B) Ergosterol Endoperoxide (EEP), and (C) Dehydroergosterol (DHE).

**Yield of sterol oxidation products after purification and quantification by HPLC**

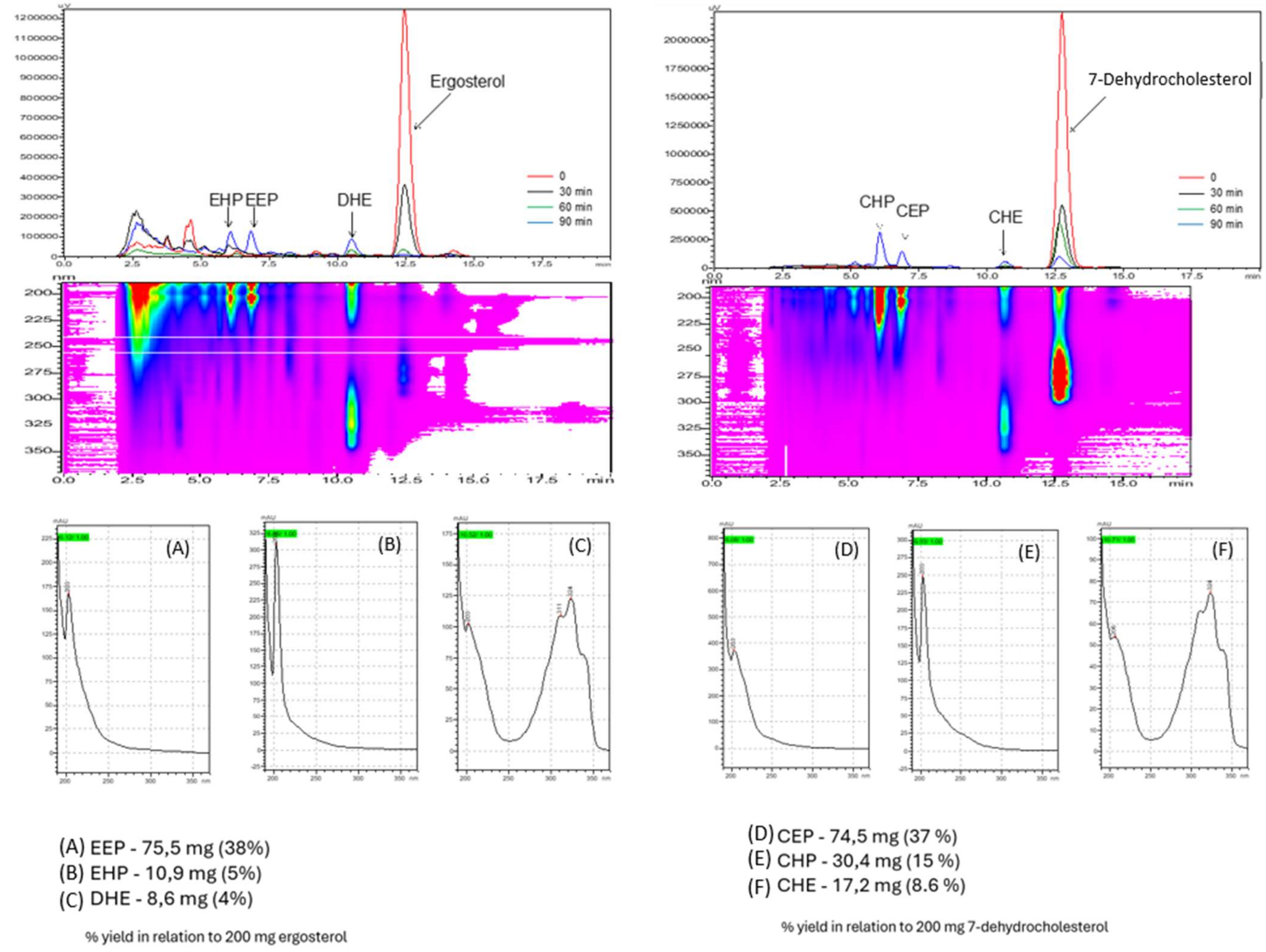


**Figure S3:** HPLC-DAD chromatograms of ergosterol and 7-dehydrocholesterol photooxidation products. The estimated yield of photo-oxidation products of sterols with methylene blue, after separation by HPLC, collecting the peaks of the respective products. Ergosterol (Erg), 7-dehydrocholesterol (7-DHC), Ergosterol Endoperoxide (EEP), Ergosterol Hydroperoxide (EHP), Dehydroergosterol (DHE), Cholesterol Endoperoxide (CEP), 8-Dehydrocholesterol Hydroperoxide (CHP), and Cholesterol Hydroperoxide (CHE).

### NMR data of ergosterol and 7-dehydrocholesterol photooxidation products

#### Ergosterol Endoperoxide (EEP)

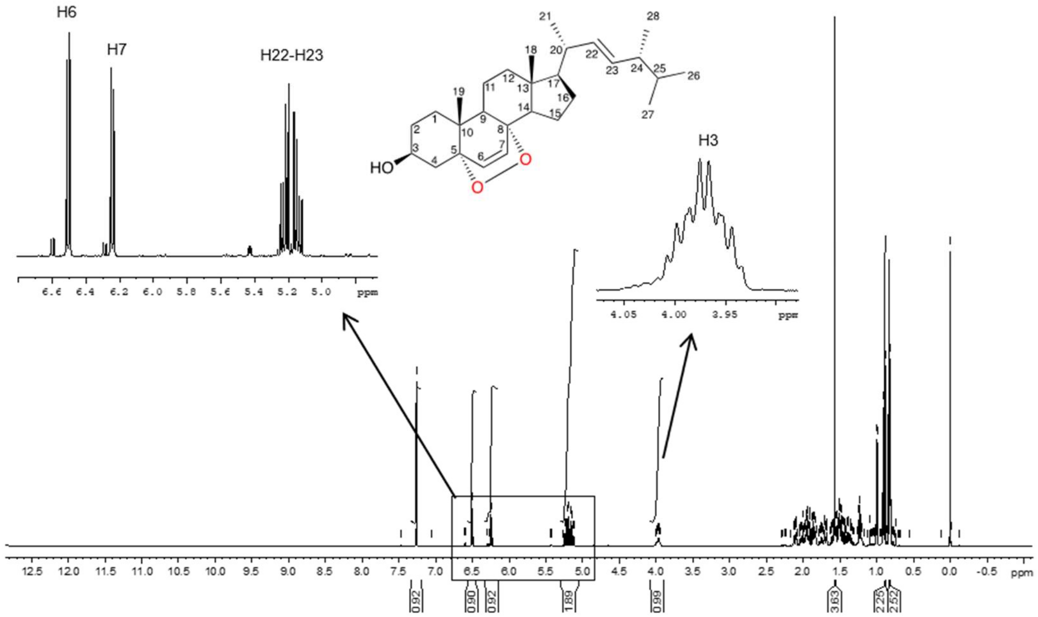


#### 7-DHC Endoperoxide (CEP)

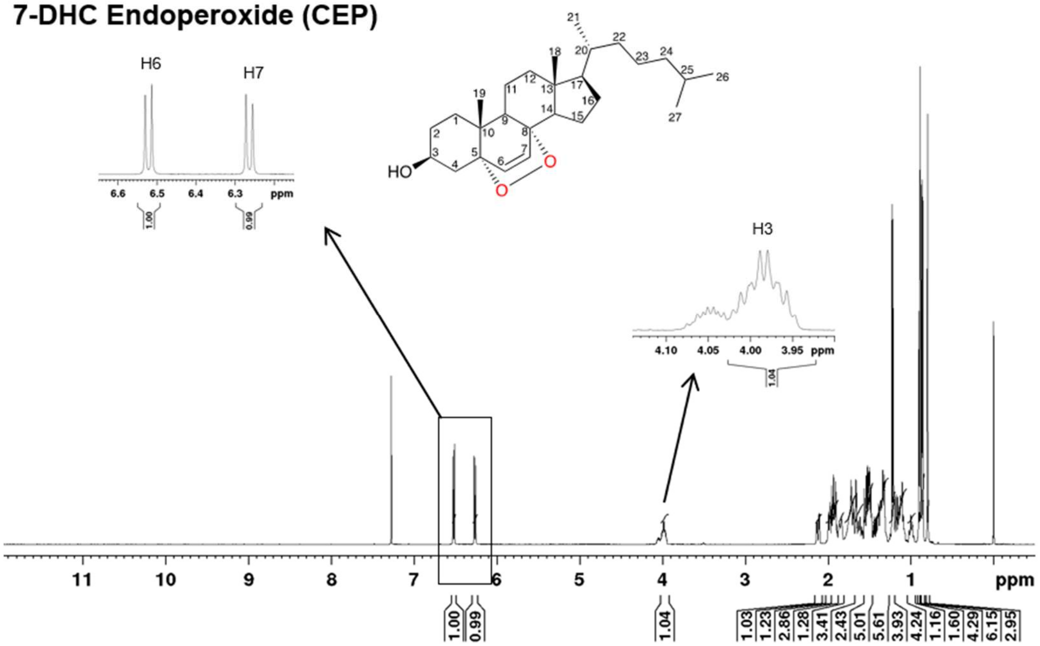


**Figure S4**. NMR spectrum of Ergosterol and 7-Dehydrocholesterol photooxidation products.

**Table S1. Chemical shifts of the main 1H observed by NMR of the fraction corresponding to EEP**

| H | **Ponce et al (ppm)** (1) | **Tian et al 2017 (ppm)** (2) | **Experimental (ppm)** |
|---|---|---|---|
| 3 | 3.85-4.03, m, J 5.3, 11.4 Hz, 1H | 3.92, m, 1H | 3.95, m, 1H |
| 6 | 6.51, d, J 8.7 Hz, 1H | 6.47, d, J 8.4 Hz, 1H | 6.52, d, J 8.5 Hz, 1H |
| 7 | 6.24, d, J 8.7 Hz, 1H | 6.21, d, J 8.4 Hz, 1H | 6.27, d, J 8.5 Hz, 1H |
| 18 | 0.82, s, 3H | 0.80, s, 3H | 0.81, s, 3H |
| 19 | 0.88, s, 3H | 0.88, s, 3H | 0.89, s, 3H |
| 22 | 5.08-5.29, m, 2H | 5.14, dd, 15.2, 8.0 Hz, 1H | 5.12, dd, 15.2, 8.5 Hz, 1H |
| 23 | 5.08-5.29, m, 2H | 5.20, dd, 15.2, 7.6 Hz, 1H | 5.21, dd, 15.2, 7.7 Hz, 1H |

**Table S2. Chemical shifts of the main 1H observed by NMR of the fraction corresponding to CEP**

| H | **Sever et al 2016 (ppm)** (3) | **Tian et al 2017 (ppm)** (2) | **Experimental (ppm)** |
|---|---|---|---|
| 3 | 3.97, m, 1H | 3.97, m, 1H | 3.98, m, 1H |
| 6 | 6.24, d, 1H | 6.51, d, 8.4 Hz, 1H | 6.26, d, J 8.5 Hz,1H |
| 7 | 6.51, d, 1H | 6.24, d, 8.4 Hz, 1H | 6.52, d, J 8.5 Hz,1H |
| 18 | 0.80, s, 3H | 0.80, s, 3H | 0.80, s, 3H |
| 19 | 0.88, s, 3H | 0.88, s, 3H | 0.88, s, 3H |
| 21 | 0.90, d, 3H | 0.90, d, 6.8 Hz, 3H | 0.89, d, J 6.6 Hz, 3H |
| 26 | 0.86, d, 3H | 0.85, d, 1.6 Hz, 3H | 0.85, d, J 2.8 Hz, 3H |
| 27 | 0.87, d, 3H | 0.87, d, 1.6 Hz, 3H | 0.87, d, J 2.8 Hz, 3H |

**Table S3. Chemical shifts (ppm) of the main 1H observed by NMR of the fraction corresponding to 7-DHC and its 7-hydroperoxide (7-OOH, CHP)**

| H | 7-DHC | 7-OOH (Experimental) | 7-OOH (Albro et al 1994) | 7-OOH (Ponce et al 2002) |
|---|---|---|---|---|
| -OOH | | 7.29, s, 1H | 7.379, s | |
| 3 | 3.66, m, 1H | 3.64, m, 1H | 3.617, m | 4.32-4.37, m, 1H |
| 6 | 5.58, dd, J 5.63 and 2.33 Hz, 1H | 5.58, dd, J 3.22 and 1.40 Hz, 1H | 5.698, d, 1.44 Hz | 5.65, dd, J 2.6 and 1.7Hz, 1H |
| 7 | 5.39, m, 1H | 4.70, ls, 1H | 4.675, bs | 5.46, br d, J 2.6 Hz, 1H |
| 18 | 0.61, s, 3H | 0.65, s, 3H | | 0.61, s, 3H |
| 19 | 0.94, s, 3H | 1.19, s, 3H | | 1.13, s, 3H |
| 21 | 0.93, d, 3H | 0.93, d, J 6.48 Hz, 3H | | |
| 26 | 0.86, d, J 2.76 Hz, 3H | 0.86, d, J 2.80 Hz, 3H | | |
| 27 | 0.87, d, J 2.76 Hz, 3H | 0.87, d, J 2.80 Hz, 3H | | |

ls= large singlet; bs=broad singlet; br d:unresolved doublet

**Di-hydroperoxide of ergosterol as photooxidation product**

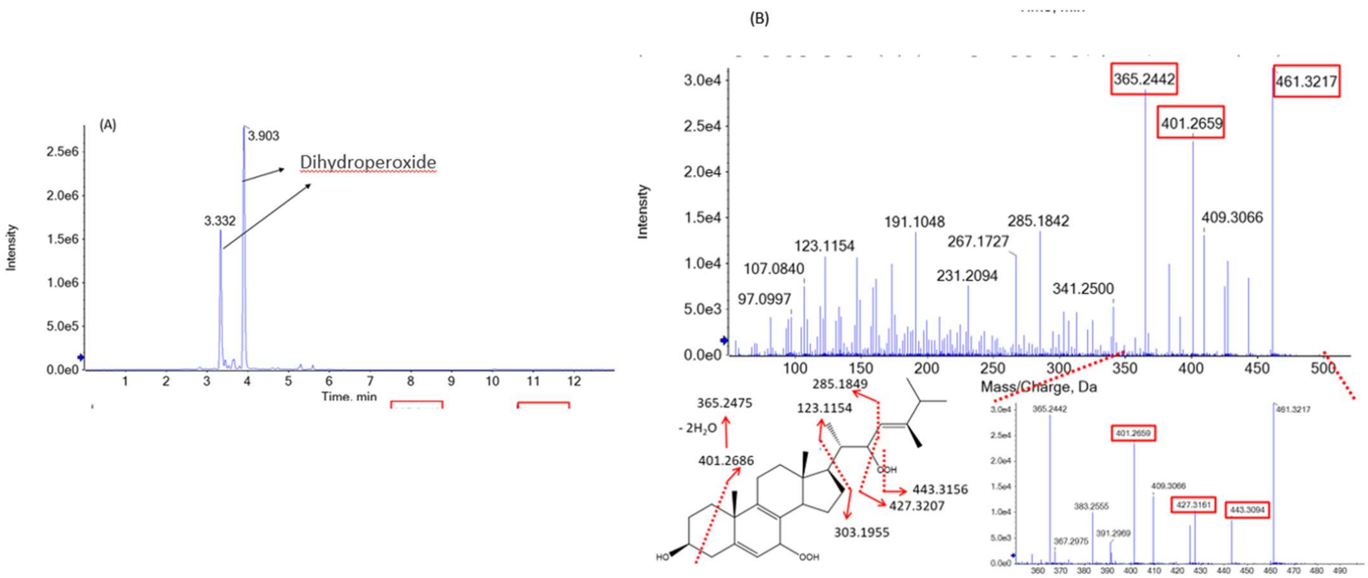


**Figure S5**. Chromatogram (A) and fragmentation spectrum (B) of ergosterol dihydroperoxide. The mass monitored was 461 m/z [M+H]+.

**Stability test of ergosterol and endoperoxides:**

Stability result showed that EHP is the least stable substance, which is expected, since hydroperoxides tend to be unstable molecules (4, 5). EEP is the most stable of all, remaining practically unchanged throughout the experiment (**Figure S6**)

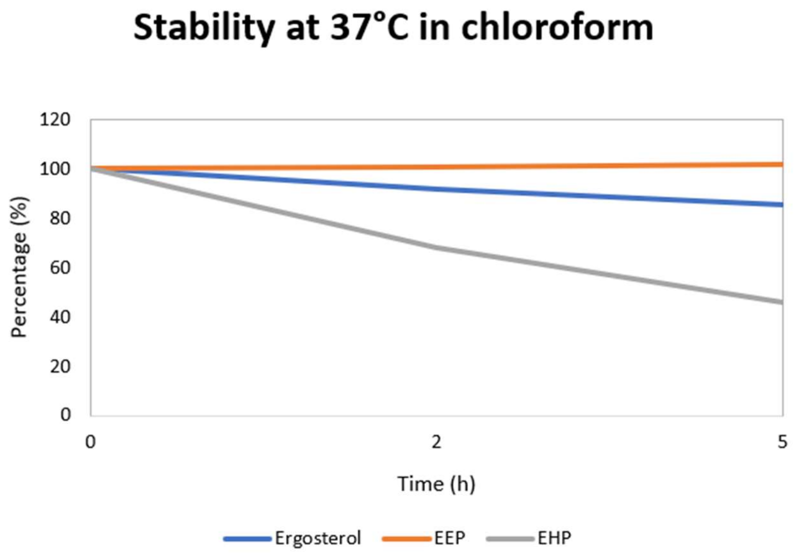


**Figure S6**. Stability of ergosterol, ergosterol endoperoxide (EEP), and ergosterol hydroperoxide (EHP) in chloroform at 37ºC stability test.

**Method of iodometry of cholesterol hydroperoxides**

Iodometry is a widely used method for quantifying hydroperoxides. The main reaction that occurs in this method is $2H^+ + ROOH + 3I^- \rightarrow H_2O + ROH + I_3^-$. This method can generate a possible interfering reaction, generating iodine at the end of the reaction, $2I^- + O_2{}^- \rightarrow I_2$.

The reaction was carried out in amber bottles and in the dark, as the presence of light can accelerate oxygen interference. Three reagents were used: acetic acid:chloroform (3:2), bubbled in $N_2$, sealed and left at room temperature; cadmium acetate (0.5 g/100 ml) also bubbled in $N_2$ in an ice bath; and potassium iodide (1.2 g/ml) which was solubilized in water bubbled with $N_2$ shortly before the experiment began.

The first stage of the experiment was the preparation of the standard curve, using the 70% tert-Butyl-OOH reagent. The concentration of the tert-Butyl-OOH stock solution was 7.77 M, with serial dilutions being made until reaching 2 mM. With this 2 mM solution, the standard curve was constructed.

To carry out the reaction with Iodine, 50 µl of the samples (standard curve or unknown sample) were placed in amber glass vials with the lights off. Then 500 µl of acetic acid: CHCl3 + 50 µl of KI were added, passing N2 for about 5s in each bottle. At the end of the process, the bottles were capped and vortexed. Each process was done separately and the entire process should take approximately 5 minutes.

At the end of this first part, 1.5 ml of cadmium acetate solution was added to stop the reaction. This solution was placed in the vials in the same order as the previous process so that all vials had the same reaction time.

After reaction, the supernatant was used to read the spectrophotometer (353 nm). 1 mol of I3 is equivalent to 1 mol of ROOH (1:1).

**Calibration curve of cholesterol and oxidation product**

The concentrations used to make the calibration curve were:

Cholesterol: 2 mM; 1 mM; 0.5 mM; 0.25 mM; 0.125 mM; 0.0625 mM

ChOOH: 1 mM; 0.5 mM; 0.25 mM; 0.125 mM; 0.0625 mM

The HPLC conditions to make the curve were: Luna C18 column (250x10 mm; 5μm, 100Å); flow of 1 ml/min; 30ºC; 30 μl injection; isocratic mobile phase with water (5%) and methanol (95%). The generated peaks were integrated, plotted on a graph and the concentration curve for each cholesterol and ChOOH were calculated.

### Evaluation of DMMB and A375 cell concentrations in photodynamic treatment

Cell viability assay with varied concentration of DMMB (0, 1, 2 and 5 nM), A375 cell concentration (3 x $10^3$, 1 x $10^4$, 5 x $10^4$ and 1 x $10^5$ cell/well) and viability measurement at 24h post-irradiation was carried out to estimate the PDT condition that minimally affects the comparative efficacy study between sterols. Following **Figure S10**, at 1 nM concentration, DMMB with PDT showed more than 70% cell viability reduction in all cell concentration. At dark condition, PDT promoted cell viability reduction of more than 50% in all cell concentration. The cell concentration also affected the viability efficacy of PDT. Thus, the combination with the sterols was carried out in a lower concentration of DMMB.

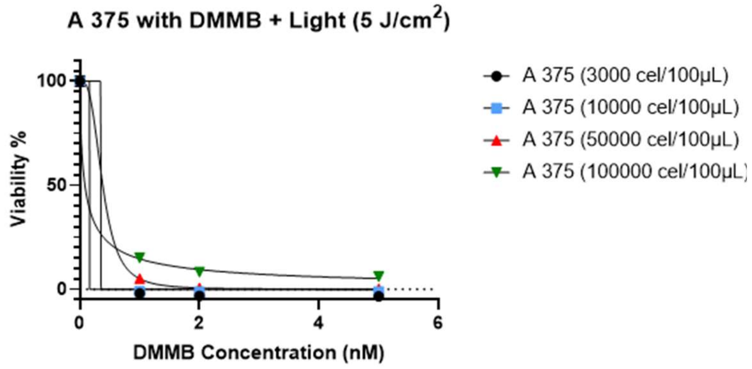


(A)

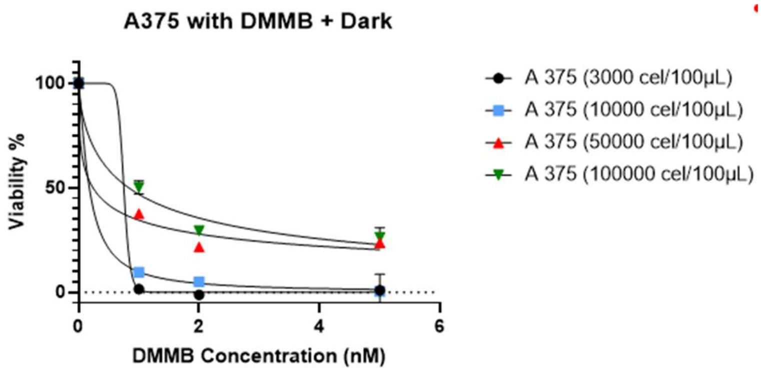


(B)

**Figure S10**. Cell viability graph of A375 with photodynamic treatment and dark condition, under different concentrations of DMMB and cell.

**Cell viability at 0 and 24 h post-irradiation**

We compared the measurement of the cell viability at 0 h and 24 h post-irradiation, with 0.1 nM DMMB, 3 x $10^4$ cell/well, and irradiation of 5 J/cm$^2$. 0h measurement demonstrated a statistically significant cell viability reduction than the 24 h measurement (**Figure S11**). Thus, the following test was conducted at 0 h measurement time.

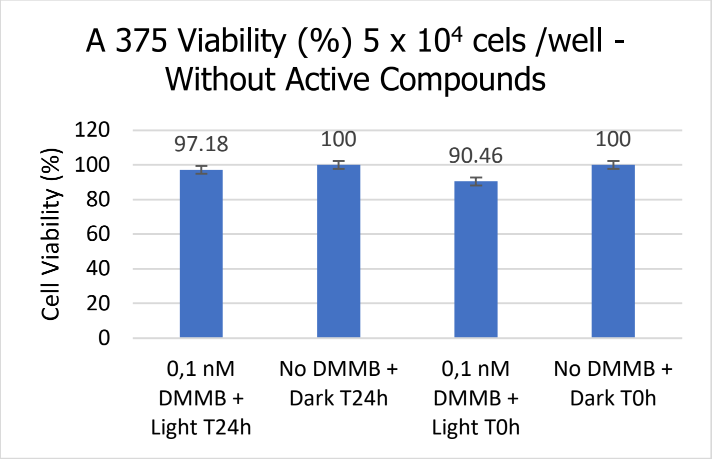


**Figure S11**. Cell viability of A 375 measurement at 0 h and 24 h post-treatment with PDT or dark condition.
